# Molecular Mechanisms Underlying Medium-Chain Free Fatty Acid-regulated Activity of the Phospholipase PlaF from *Pseudomonas aeruginosa*

**DOI:** 10.1101/2023.05.02.539057

**Authors:** Rocco Gentile, Matea Modric, Björn Thiele, Karl-Erich Jaeger, Filip Kovacic, Stephan Schott-Verdugo, Holger Gohlke

## Abstract

PlaF is a membrane-bound phospholipase A_1_ from *P. aeruginosa* that is involved in remodeling membrane glycerophospholipids (GPLs) and modulation of virulence-associated signaling and metabolic pathways. Previously, we identified the role of medium-chain free fatty acids (FFA) in inhibiting PlaF activity and promoting homodimerization, yet the underlying molecular mechanism remained elusive. Here, we used unbiased and biased molecular dynamics simulations and free energy computations to assess how PlaF interacts with FFAs localized in the water milieu surrounding the bilayer or within the bilayer, and how these interactions regulate PlaF activity. Medium-chain FFAs localized in the upper bilayer leaflet can stabilize inactive dimeric PlaF, likely through interactions with charged surface residues as experimentally validated. Potential of mean force (PMF) computations indicate that membrane-bound FFAs may facilitate the activation of monomeric PlaF by lowering the activation barrier of changing into a tilted, active configuration. We estimated that the coupled equilibria of PlaF monomerization-dimerization and tilting at the physiological concentration of PlaF lead to the majority of PlaF forming inactive dimers when in a cell membrane loaded with decanoic acid (C10). This is in agreement with a suggested *in vivo* product feedback loop and GC-MS profiling results indicating that PlaF catalyzes the release of C10 from *P. aeruginosa* membranes. Additionally, we found that C10 in the water milieu can access the catalytic site of active monomeric PlaF, contributing to the competitive component of C10-mediated PlaF inhibition. Our study provides mechanistic insights into how medium-chain FFA may regulate the activity of PlaF, a potential bacterial drug target.

## 1. INTRODUCTION

The Gram-negative bacterium *P. aeruginosa* is a human pathogen and a frequent cause of nosocomial infections, affecting primarily immune-compromised patients (1, 2). A large spectrum of virulence factors contributes to the pathogenicity of *P. aeruginosa* (3). Among these, type A phospholipases (PLA_1_) participate in host membrane damage and modulation of various signaling networks in infected cells (4, 5). PLA_1_ hydrolyzes membrane GPL at the *sn*-1 position yielding lysoglycerophospholipid (LGPL) and FFA (6, 7). GPLs are membrane components involved in cellular integrity and regulation of membrane protein function and stability (8). The alteration of the membrane GPL composition has been linked to biofilm formation, growth phase transition, virulence, and cytotoxicity of *P. aeruginosa* (1, 2, 6, 9). On the other side, FFA in *P. aeruginosa* acts as a signaling molecule (10) or as the precursor of signal molecules including the hydroxy-alkylquinolines (11), diffusible signal factors or oxylipin autoinducer (12) families, important for bacterial virulence in the pathogen (13).

Previously, we showed that *P. aeruginosa* PlaF is a cytoplasmic membrane-bound PLA_1_ that contributes to the alteration of the membrane GPL profile and virulence properties of this bacterium (6). *In vitro* enzyme activity experiments showed a specificity of PlaF towards GPLs and LGPLs with medium-chain acyl moieties (10 - 14 carbon atoms), while short- and long-chain GPLs are only poorly hydrolyzed (6, 15). A high-resolution crystal structure of PlaF, combined with homodimerization studies by crosslinking, micro-scale thermophoresis, and molecular simulations, has revealed that PlaF can adopt both monomeric and dimeric configurations **(Figure 1)** (6). However, only the monomeric state of the protein was found to be catalytically active. The crystal structure revealed a homodimer characterized by interactions between the transmembrane (TM) and juxtamembrane (JM) regions between the single monomers. The crystal structure of the PlaF dimer revealed the co-crystallized endogenous ligands undecanoic acid (11A) and tetradecanoic acid (myristic acid, MYR) in the catalytic site of each monomer, with their acyl chains occupying tunnel 1 (T1) pointing towards the dimeric interface (6). Free energy computations (16) and experiments indicated that, at physiological protein concentrations, the equilibrium between the PlaF dimer and the monomers is shifted to the latter side in the cell (6). In the dimeric form, the entrance of T1 resides more than 5 Å above the membrane’s upper leaflet, thus hampering GPL substrate access from the membrane in T1 (6). Interestingly, unbiased molecular dynamics (MD) simulations (17) showed that the PlaF monomer undergoes a tilting motion in the membrane, bringing the opening of the catalytic tunnel in close vicinity to the membrane, with free energy computations revealing that the tilted state of the monomeric PlaF (t-PlaF) is energetically more favorable than the non-tilted state, named split or s-PlaF state (**Figure 1**) (6). Consequently, it has been suggested that this tilting motion underlies the activation of the monomeric s-PlaF by facilitating substrate access to the active site tunnel (6).

**Figure 1.**
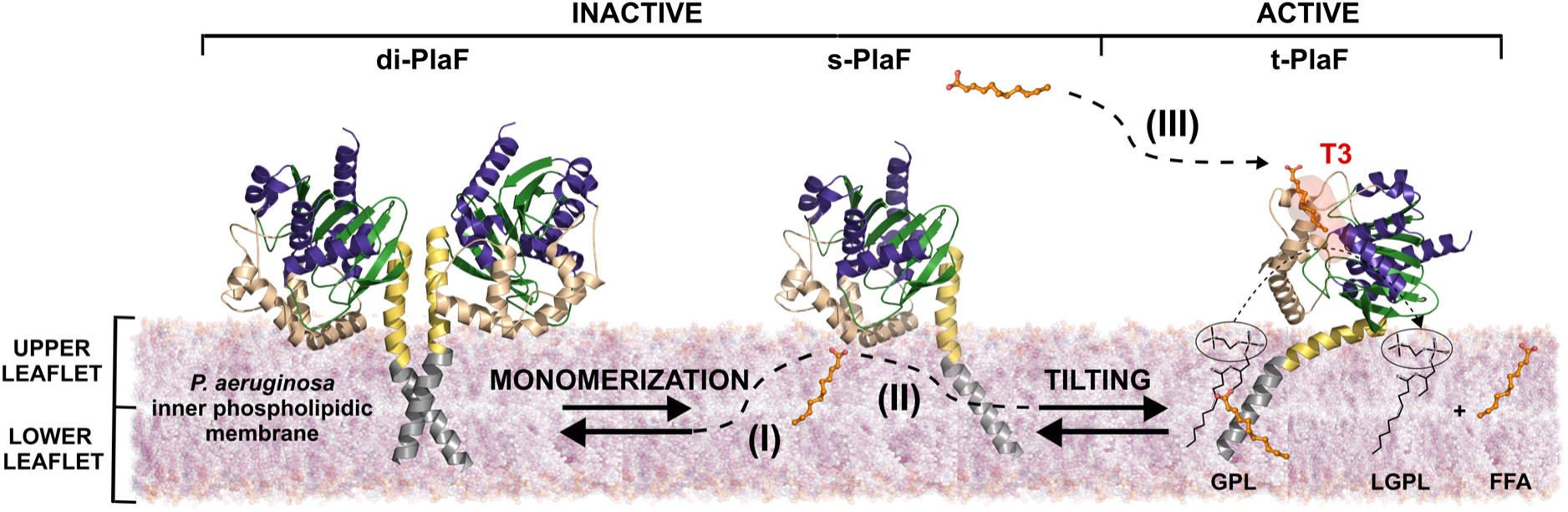
A working model illustrating the possible impact of FFA on the dimer-monomer equilibrium of PlaF and its monomer tilting. PlaF is anchored to the bilayer through a transmembrane (TM) region (grey) predominantly containing hydrophobic residues and a juxtamembrane (JM) region (yellow) rich in polar and charged residues. The JM domain is in close contact with the polar heads of the phospholipid membrane when PlaF is tilted. The C-terminal catalytic domain points towards the periplasm and is characterized by an α/β hydrolase fold (green, violet) providing the scaffold for the catalytic triad (Ser, Asp, His) and Arg-Lys-rich lid-like (LL, light brown) domain that presumably interacts with the phospholipid bilayer. PlaF can adopt a dimeric (di-PlaF) or monomeric configuration (s-PlaF), and the monomer can tilt (t-PlaF). The t-PlaF monomer is considered the active form (6). Lipid bilayer-bound FFA (orange), generated by PlaF from GPL or LGPL, may influence the dimer-monomer equilibrium (I) and/or the tilting transition (II) and/or enter PlaF from the water milieu (III) through tunnel 3 (T3).

In addition, two other tunnels (T2 and T3) that connect the active site to the surface of PlaF have been identified (15). Unbiased MD simulations suggested that MYR, the hydrolysis product of GPL and LGPL, relocates from T3 and reaches the entrance of T1, in agreement with the positions of cocrystallized ligands MYR and 11A (6, 15). Accordingly, the FFA can reach the membrane bilayer *via* T1. We have recently shown that medium-chain FFAs, containing 10 - 14 carbon atoms, inhibit PlaF activity according to a mixed inhibition mode and that they lead to an increase in the PlaF dimer concentration (6). From our results, we hypothesize that the FFAs may inhibit PlaF activity by two mechanisms: i) allosterically, by impacting the dimer-monomer equilibrium and/or the tilting transition when they are located in the lipid bilayer, and ii) competitively, by interfering with the substrate binding when they enter the active site *via* T3 from the water milieu mimicking the periplasmic space **(Figure 1)**. However, the detailed molecular mechanisms that govern the PlaF inhibition by FFAs have remained elusive.

Here, we combine unbiased and biased all-atom MD simulations (18) and configurational free energy computations (16) to evaluate the effect of FFAs on the structural dynamics and energetics of the PlaF dimer-monomer transition and the monomer tilting and to reveal hot spot PlaF residues potentially involved in the interaction with FFAs. Additionally, we pursue free ligand diffusion MD simulations (19) and binding free energy computations (16) of FFAs located in the water phase near the catalytic domain of PlaF. The suggested FFA binding sites in PlaF and their putative roles in inhibition and dimerization were experimentally studied using site-directed mutagenesis and purified PlaF variants reconstituted into small unilamellar vesicles (SUVs) in inhibition and crosslinking assays.

## 2. RESULTS

### 2.1 Medium-chain Fatty Acids are Detected in *P. aeruginosa* in Free Form and Likely Originate from Glycerophospholipid Hydrolysis

Our previous lipidomics analysis by Q-TOF MS/MS revealed that the alteration of the membrane GPL profile of *P. aeruginosa* is linked to PlaF-mediated degradation of several GPLs, including three GPLs containing medium-chain FAs: phosphatidylglycerol (PG) 24:3, phosphatidylethanolamine (PE) 22:1, and phosphatidylinositol (PI) 26:0 (6). In this study, and to evaluate how PlaF modifies the FFA profile and the FFA availability to potentially regulate PlaF activity in the periplasmic space, we analyzed if medium-chain FFAs can be identified in *P. aeruginosa* wild-type (WT) and an isogenic *plaF* mutant (Δ*plaF*) with reduced GPL degrading activity (6). To achieve this, we extracted FFAs from cells and supernatant of both strains using organic solvent and quantified the saturated FFAs with acyl chains of 6 to 16 carbon atoms by GC-MS. The FFA profile revealed that both *P. aeruginosa* strains produced various intracellularly and extracellularly located saturated FFAs **(Figure 2)**. As anticipated, the long-chain FFAs were the most abundant, while medium-chain FFAs (C8, C9, C10, C12, and C14) accounted for roughly 10 to 15 % of the total FFA amount. No significant differences in the abundance of cell-associated FFAs between the WT and Δ*plaF* were observed, suggesting that FFAs released by PlaF are either metabolized or secreted into the supernatant. We identified that medium-chain FFAs, namely octanoic (C8), decanoic (C10), dodecanoic (C12), and myristic (C14) acids, were significantly less abundant (*p* < 0.05) in the supernatant of Δ*plaF* than the WT.

**Figure 2:**
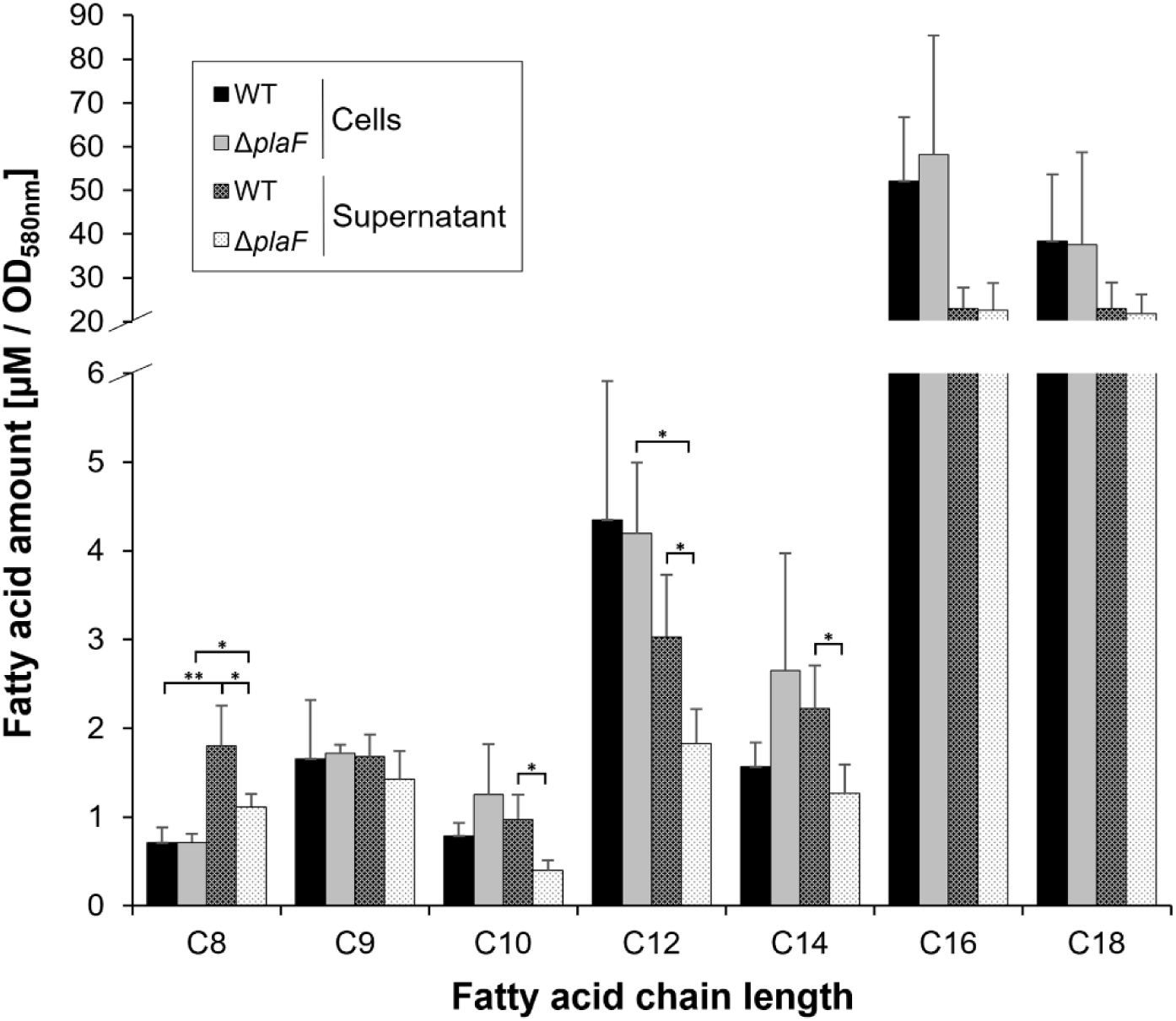
Free fatty acid profile of *P. aeruginosa* WT and Δ*plaF*. Strains were cultivated in LB medium under aeration until the stationary phase, and FFAs were extracted with organic solvent from cells and supernatant. Quantification of FFAs was performed by GC-MS analysis. Means ± SDs are shown (*n* = 4 biological replicates). Statistical analysis was performed using the *t*-test, * *p* < 0.05, ** *p* < 0.01; differences between all other data points were not significant.

To further explore the possibility that PlaF releases the identified medium-chain FAs from GPLs, we conducted a re-analysis of our published lipidomics results obtained with the Δ*plaF* mutant (6). Our previous analysis had assessed GPLs at the species level, meaning that only information about the total number of carbon atoms in the acyl chains but not their specific identities had been obtained. Thus, we focused on GPL species with up to 24 carbon atoms in their acyl chains, as they likely contain at least one medium-length acyl chain (**Table S1**). Our results show that the *P. aeruginosa* membrane contains 56 different medium-chain length GPLs out of a total of 324 identified GPL species (**Table S1**). A comparison of the amount of medium-chain length GPLs relative to the total GPL amount in WT and Δ*plaF* revealed that more (20.8 %) medium-chain length GPLs were identified in the Δ*plaF* mutant than the WT (12.6 %) (**Table S2**). These results together with the specificity of PlaF to hydrolyze medium-chain GPLs (6) suggest that the identified medium-chain FFAs could potentially be generated by PlaF-catalyzed hydrolysis of GPLs.

The GC-MS analysis revealed an interesting finding: the extracellular octanoic acid amount was nearly double (*p* < 0.001) that of the intracellular amount in both the WT and Δ*plaF* strains, suggesting that this FFA is predominantly secreted into the supernatant. In contrast, dodecanoic acid, the second FFA identified as a potential PlaF product, was significantly (*p* < 0.001) more abundant inside the cells of Δ*plaF* than the WT. These results show that the FFA transport is dependent on their chain length, as it was observed previously for *Escherichia coli* and *Saccharomyces cerevisiae* (20). This suggests that some FFAs may have a function as extracellularly secreted compounds, such as signaling messengers (10), while others may be primarily used intracellularly, such as for energy production (21).

To conclude, the quantification of intracellular and extracellular FFAs is in agreement with our previously determined ability of PlaF to release FFA *in vitro* from various GPLs (6), and further strengthens the suggested *in vivo* function of PlaF in degrading GPLs containing medium-length acyl chains.

### 2.2 Free Fatty Acids in the Membrane may Facilitate the Tilting of PlaF Monomers

To assess the influence of bilayer-bound FFA on PlaF tilting, we performed all-atom unbiased MD simulations (18) of monomeric PlaF inserted into the lipid bilayer loaded with FFA. The orientations of PlaF obtained after splitting the membrane-embedded PlaF dimer (s-PlaF) or the tilted configuration (t-PlaF) obtained with OPM were considered (22, 23). For each PlaF configuration, two membrane systems were examined, in which C10 or C14 were inserted in the upper leaflet resulting in a DOPE:DOPG:FFA composition of 3:1:1. This system mimics the FFAs bound to the periplasmic leaflet of the inner bacterial membrane, which is interacting with the TM, JM and KR-rich LL domains of PlaF (6). For each system, 12 replicas were simulated, each 1 μs in length, and the results were compared with those obtained previously with PlaF in an FFA-free bilayer composed of DOPE and DOPG at a ratio of 3:1, respectively (6).

When starting from the s-PlaF orientation, a comparable number of transitions to the t-PlaF orientation (“tilting”) was observed for the system containing C10 (9/12, 75%) or C14 (10/12, 83%) **(Figure 3)**. These tilting preferences are higher compared to the 50% obtained previously for the system without FFAs (6). These results indicate that FFAs may facilitate the tilting and that for the tested FFA chain length has no influence on the tilting preference. When starting MD simulations from the t-PlaF embedded into the C10- or C14-loaded bilayer, no transition to the s-PlaF orientation was observed **(Figure 3)**, even if the MD simulations were prolonged to 2 μs **(Figure S1)**. These results are in agreement with our previous finding that the tilted orientation of PlaF is preferred (6).

### 2.3 Free Fatty Acids Favor Monomer Tilting but Disfavor Dimer Dissociation

To determine the effect of FFAs in the upper leaflet on the energetics of the PlaF dimer-to-monomer and s-PlaF-to-t-PlaF transitions, we performed umbrella sampling (US) simulations (24) to compute a PMF (25). As the chain length showed a negligible influence on the tilting behavior in the unbiased MD simulations, we only used a membrane system with DOPE:DOPG:C10 of 3:1:1. The US simulations were performed as in ref. (6), starting from the s-PlaF orientation. The distance between the top of the JM domain, computed as the center of mass (COM) of C_α_ atoms of residues 33-37, and the membrane center along the membrane normal was used as a reaction coordinate to track the tilting transition. For the dimer-to-monomer transition, we started from the dimer configuration as found in the crystal structure and used the distance between the COMs of C_α_ atoms of residues 25-38 of each PlaF molecule as a reaction coordinate. For the tilting transition, 43 windows were sampled with 800 ns of sampling time each, of which the first 640 ns were discarded as equilibration. Likewise, for the dimer-to-monomer transition, 75 windows were sampled with 800 ns of sampling time each, of which the first 640 ns were discarded as equilibration. In both cases, the kernel densities showed a median overlap of 34.9 ± 1.4% and 32.2 ± 1.5% between contiguous windows, respectively **(Figures S7 and S8)**, well suited for PMF calculations (26). The standard error of the mean was calculated from free energy profiles determined independently every 20 ns from the last 160 ns per window. The PMF of the titling transition is converged and precise **(Figure S7)**. In the presence of C10 in the bilayer, t-PlaF is preferred over s-PlaF **(Figure 4A)**, which is similar to results without FFA in the bilayer (6). However, the global minimum of t-PlaF is lower by ∼4 kcal mol^-1^ in the bilayer containing C10 than the one without C10, indicating that t-PlaF is even more favorable in the presence of C10. Additionally, a local minimum was observed in the system containing C10 instead of a barrier separating the split and tilted PlaF configurations in the system without FFA **(Figure 4A)**. These results are in accordance with the unbiased MD simulations, which indicated that the tilted configuration is preferred and that the transition to it is facilitated in the presence of FFAs.

**Figure 3.**
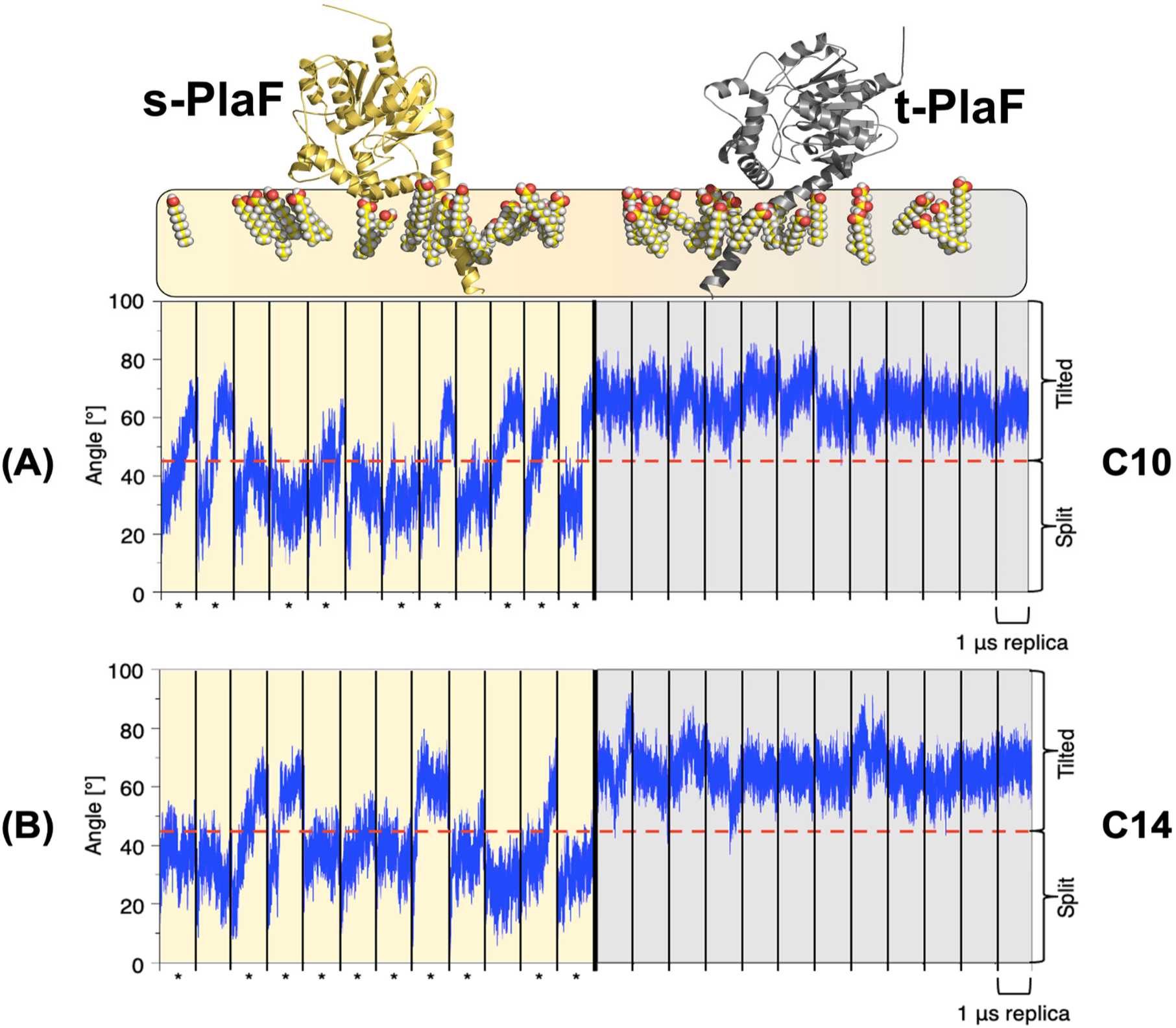
Unbiased MD simulations of monomeric PlaF in the presence of FFA in the upper leaflet. **(A)** Left (yellow background): Time course of the orientation of PlaF with respect to the bilayer starting from s-PlaF in the presence of C10 at a ratio DOPE:DOPG:C10 of 3:1:1. In 9 of 12 replicas, the s-PlaF adopts a tilted configuration marked with *. Right (grey background): When starting from t-PlaF, the structure remains tilted in all simulations. As before (6), this shows a significant tendency of the monomer to tilt (McNemar’s χ^2^ = 6.125, *p* = 0.013). Tilting is quantified by the angle between the membrane normal and the vector between the center of masses (COM) of C_α_ atoms of residues 21–25 and residues 35–38 and is assumed to occur if the angle is > 45°. **(B)** As in panel A, but now with C14 at a ratio of DOPE:DOPG:C14 of 3:1:1. In 10 of 12 replicas, s-PlaF adopted a tilted configuration (marked with *, McNemar’s χ^2^ = 8.100, *p* = 0.004) while t-PlaF did not undergo a transition to s-PlaF.

**Figure 4.**
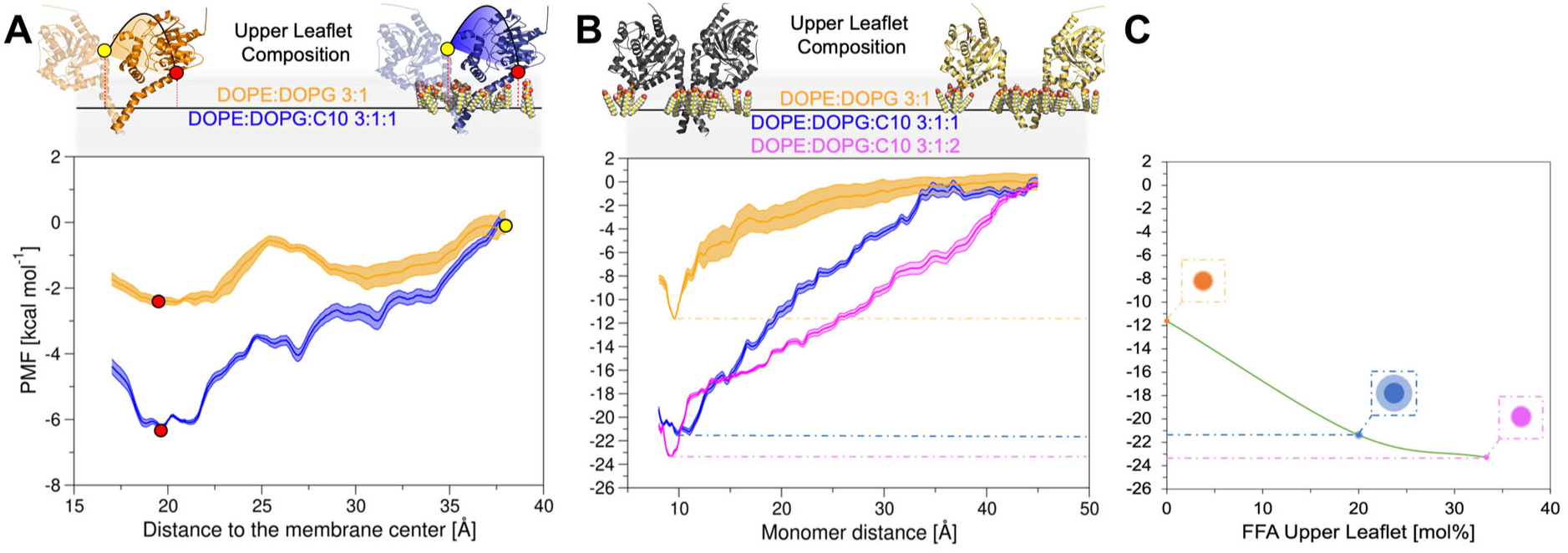
PMFs of monomer tilting and dimer separation. **(A)** PMF of monomer tilting. The distance along the membrane normal between the COM of C_α_ atoms of residues 33-37 and the COM of the C18 of the oleic acid moieties of all GPLs in the bilayer was used as a reaction coordinate. Orange: PMF computed for the bilayer consisting of DOPE:DOPG 3:1 (values were taken from ref. (6)). Blue: PMF computed here for the bilayer containing C10 in the upper leaflet at the ratio DOPE:DOPG:C10 of 3:1:1. The shaded areas at the curves show the standard error of the mean (SEM). The yellow and red dots indicate s-PlaF and t-PlaF, respectively. **(B)** PMF of di-PlaF separation. The distance between the COM of C_α_ atoms of residues 25-38 of each PlaF molecule was used as the reaction coordinate. Orange: PMF values for the bilayer consisting of DOPE:DOPG 3:1 were taken from ref. (6)). Blue: PMF for the bilayer containing C10 in the upper leaflet at the ratio DOPE:DOPG:C10 of 3:1:1. Magenta: PMF computed for the bilayer containing C10 in the upper leaflet at the ratio DOPE:DOPG:C10 of 3:1:2. The shaded areas show the SEM. The black protein on the top shows the PlaF dimer at a monomer distance of ∼9.9 Å, and the yellow ones show the completely dissociated PlaF at ∼45 Å. **(C)** PMF values at the respective global minima. This panel shows PMF values at the respective global minima from panel B as a function of the C10 concentration in the upper bilayer leaflet. Dashed lines connect respective points in panels B and C. The green line connecting the points illustrates a non-linear correlation between FFA concentration in the upper leaflet and PlaF dimer stabilization. The shaded area around each dot indicates the SEM at the corresponding minimum.

The PMF for the dimer-to-monomer transition was computed considering two bilayer compositions with 20 mol% and 33.3 mol% of C10 in the upper leaflet. The PMFs are converged and precise **(Figures S8 and S9)**. As in the membrane without FFA (6), dimeric PlaF is favored over the monomeric state in the presence of C10 in the membrane **(Figure 4B)**. However, FFA stabilizes the di-PlaF in a concentration-dependent manner by ∼10 and ∼14 kcal mol^-1^ compared to a system without FFA.

To conclude, the PMF computations demonstrated that the presence of C10 within the upper bilayer leaflet promotes the tilting of s-PlaF, corroborating the findings of our MD simulations (**Figure 3A**), and C10 exhibits a concentration-dependent stabilizing effect on the PlaF dimer **(Figure 4C)**.

### 2.4 Estimating the Ratio of Monomeric and Dimeric PlaF in the Cell in the Presence of Free Fatty Acids

Following previous work by us (6, 27) and others (28), we computed from the PMFs of dimer-to-monomer and tilting transitions with an upper leaflet composition of DOPE:DOPG:C10 of 3:1:1 equilibrium constants (*K_a_* = 1.68 × 10^14^ Å^2^ (eq. S1), *K_x_* = 2.59 × 10^12^ (eq. S2), *K_tilting_* = 1.73 × 10^2^ (eq. S5)) and free energies (Δ*G* = - 17.04 ± 0.22 kcal mol^-1^ (eq. S3), Δ*G_tilting_* = - 3.04 ± 0.20 kcal mol^-1^ (eq. S6)) taking into account that *K_x_* and Δ*G* relate to a state of one PlaF dimer in a membrane of 1258 GPLs, according to our simulations setup. Experimentally, a concentration of one PlaF dimer per ∼3786 GPLs in *P. aeruginosa* PlaF-overexpressing cells was determined (29). However, the concentration in *P. aeruginosa* wild-type (WT) is estimated as 100- to 1000-fold lower.

Under such physiological conditions in the *P. aeruginosa* WT and considering that the equilibria for dimer-to-monomer transition and tilting are coupled, between 2.3 and 7.3% of the PlaF molecules are predicted to be in a monomeric, tilted, catalytically active state in *P. aeruginosa* when 20 mol% C10 are present in the upper leaflet. In the absence of FFAs, between 74% and 96% of the PlaF molecules were predicted to be in a monomeric, tilted, catalytically active state in *P. aeruginosa* WT (6). *Vice versa*, in the presence of FFA, 92.7 to 97.7% of PlaF are in the dimeric configuration, whereas this is only 4 to 26% in the absence of FFA (6). The computed increase in the PlaF dimer concentration agrees with our previous biochemical results showing an increased di-PlaF concentration after incubating purified PlaF with C10 (6). Based on these findings, as well as the GC-MS results **(Figure 2)** that indicate the release of C10 by PlaF *in vivo*, it is plausible to propose that the degradation of GPLs catalyzed by PlaF is inhibited by the produced C10. This inhibition may serve as a protective product-feedback mechanism to prevent cell membrane damage.

### 2.5 Hot spots of di-PlaF – FFA Interactions in the Upper Leaflet Reduce Experimental Dimer Stability when Mutated

To characterize interactions between C10 in the upper leaflet and di-PlaF that may favor dimer formation, we analyzed the trajectories of unbiased MD simulations across 12 replicas. 3D density grids representing the probability density of C10 within the membrane led to the identification of states in which C10 is located at a distance of ≤ 5 Å from the PlaF. To discard short-living di-PlaF-C10 states we analyzed if the RMSD of C10 in two consecutive states is < 1.5 Å, indicating that these states encompassed C10 tightly bound to di-PlaF (see Supplementary results). They were clustered with respect to the minimum distance *ε* between the clusters starting from *ε* = 2.0 Å, using the all-atom RMSD as the similarity measure. Gradual increase of *ε* in 0.5 Å intervals led to a constant population of the largest cluster at *ε* = 5.0 Å.

The five most populated clusters cover 80.5 ± 1.5% (mean ± SEM) of all systems with C10 close to PlaF and 65.8 ± 1.3% of all systems in a trajectory **(Figure 5A)**. Over these clusters, R187 and R217, with positively charged side chains, E216 containing a negatively charged side chain, and A220 with a non-polar side chain were identified as the residues mostly interacting with FFA **(Figure 5B)**. These residues are surface-exposed and located in the LL domain, which according to the model of di-PlaF in the membrane interacts with the bilayer (6).

**Figure 5.**
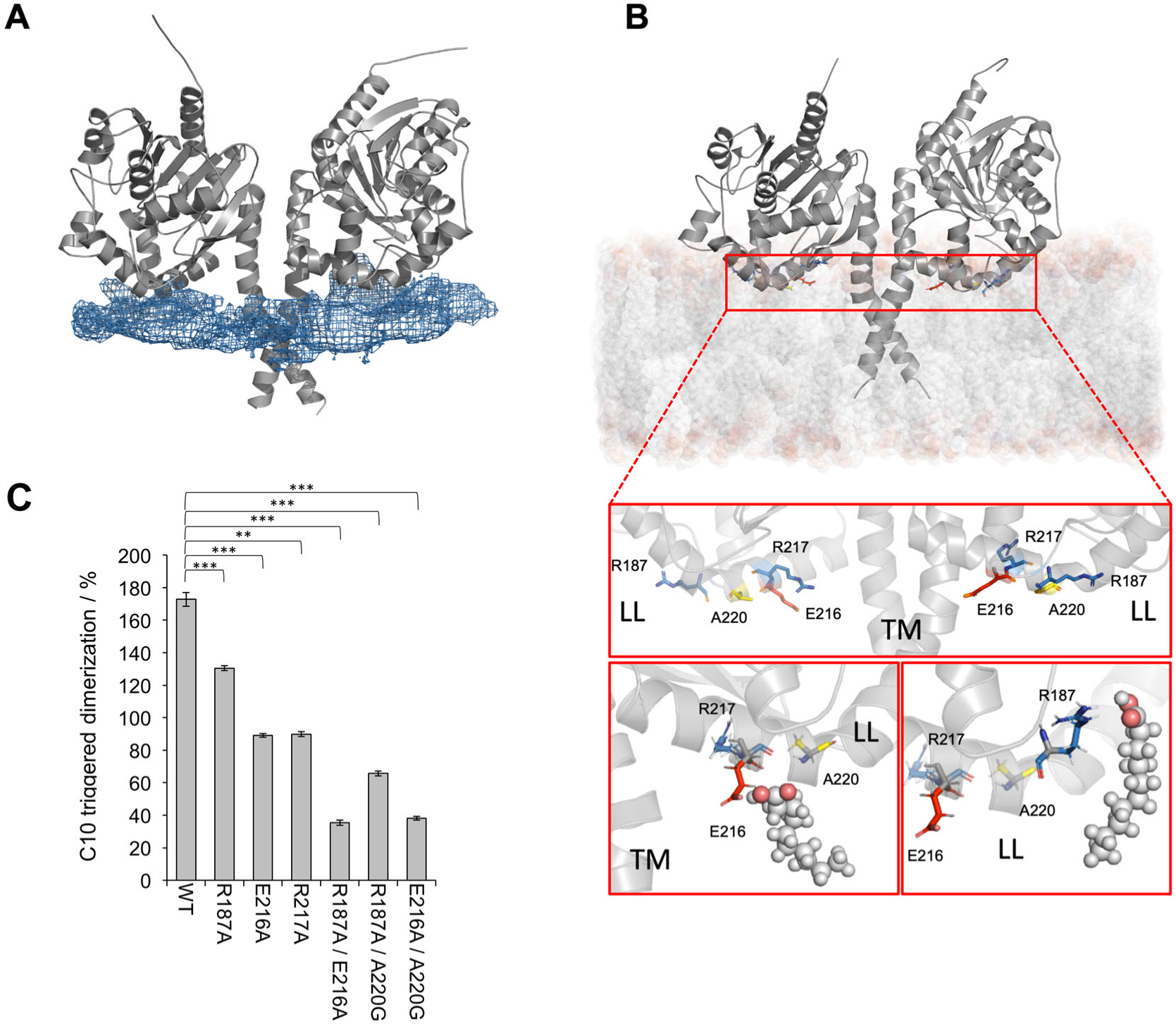
Interaction of C10 located within the upper leaflet with di-PlaF. **(A)** Average density map of all C10 molecules. 3D density grid illustrating the distribution of all C10 computed with CPPTRAJ (33). The C10 distribute around both monomers in similar proportions. **(B)** Hot spots of di-PlaF-C10 interactions. Hot spots were defined as residues with a distance to C10 ≤ 5 Å and where C10 has an RMSD < 1.5 Å with respect to the previous pose. The red box contains the negatively and positively charged hot spots, shown in red and blue sticks, respectively, while non-polar ones are reported in yellow sticks; LL, the lid-like domain; TM, the transmembrane domain. Red frames at the bottom show enlarged interaction regions with C10 shown in space-fill format (grey, carbon; red, oxygen). The left panel indicates the interaction of C10 with E216 and R217. The right panel indicates the interaction of C10 with R187. **(C)** Effect of mutations in the LL domain on PlaF dimerization. PlaF_WT_ or PlaF variants reconstituted into DOPE:DOPG SUVs were treated with C10 (12 mM) and crosslinker DMP for 2 h at room temperature. SUV reconstituted proteins treated with DMP and DMSO (used as C10 solvent) served as the control without C10. Proteins were separated by SDS-PAGE, followed by Western blot detection of PlaF using anti-His-tag antibodies. The intensities of di-PlaF bands were quantified by ImageJ (34). The results represent the ratio between C10-untreated and C10-treated samples shown as mean ± SD (*n* = 4).

To experimentally test the role of these residues on PlaF dimerization in the GPL bilayer, we mutated by site-directed mutagenesis each of the three charged residues to alanine to obtain the single variants PlaF_R187A_, PlaF_E216A_, and PlaF_R217A_. Furthermore, a variant with two neutralized charges, PlaF_R187A-E216A_, and variants with one neutralized charge and C10-interacting A220 mutated to glycine, PlaF_R187A-A220G_ and PlaF_E216A-A220G_, were generated. These six PlaF variants and the wild-type PlaF (PlaF_WT_) were produced in *E. coli* and purified by immobilized metal affinity chromatography (IMAC) in the presence of octyl β-D-glucopyranoside (OG). PlaF_WT_ and variants purified to homogeneity as determined by SDS-PAGE **(Figure S10A)** showed comparable thermal stability as determined by nano differential scanning fluorimetry (nanoDSF) **(Table S4)**. These results indicate that the mutations did not markedly destabilize the protein, as mutations of surface-exposed residues are on average considerably less destabilizing compared to mutations of core residue positions (30).

Furthermore, we removed OG **(Figure S10B)** and reconstituted PlaF_WT_ and variants into SUVs made of DOPE and DOPG, and C10 was added to reach 20% mol. In the experimental setup, the pH was 8, surpassing the p*K*a value of C10 (6.4), such that C10 is predominantly in a deprotonated and non-micellized state (31). In this state, C10 has limited permeability through the phospholipid bilayer, consequently favoring its partitioning within the outer leaflet (32). This experimental system closely approximates the computational model, rendering it suitable for subsequent comparison with computational data.

A quantitative crosslinking assay **(Figure S11)** using the bifunctional reagent dimethyl pimelimidate (DMP), which covalently stabilizes dimers, revealed that C10 triggers the formation of PlaF_WT_ dimers **(Figure 5C)**, as shown before (6). We observed 70% more PlaF_WT_ dimer in the C10-treated sample compared to the untreated one. In contrast, all PlaF variants showed a significantly reduced ability to dimerize in the presence of C10. For single variants, relative amounts of di-PlaF in C10-treated samples were approximatively equal compared to untreated samples. In double-point variants, less di-PlaF in C10-treated than C10-untreated samples was observed. These results indicate that residues predicted by MD simulations to interact with C10 within the bilayer play an important role in C10-triggered PlaF dimerization.

### 2.6 Free Fatty Acids in the Aqueous Milieu Interact with t-PlaF Primarily via Entering T3

Subsequently, we investigated if C10 from the aqueous milieu could directly bind to PlaF without prior entry into the bilayer. To explore this, we conducted all-atom unbiased MD simulations of free ligand diffusion (fldMD) of 1 µs length. Such simulations have been previously used to predict the binding modes of molecules (19). t-PlaF was embedded in a bilayer with the composition DOPE:DOPG of 3:1 and 10 molecules of C10 were added to the water phase, resulting in concentrations of 30 mM in the solvation box. This higher concentration than the experimentally used one was chosen to increase the probability of observing events of C10 binding to and dissociating from t-PlaF. As this concentration is above the CMC of C10 (35), we monitored by visual inspection of the trajectories that no C10 aggregates formed.

We performed 12 replicas, and binding and unbinding events of C10 were monitored **(Figure S12)**. We identified a C10 as “bound” if it is within a distance < 5 Å from the T3 entrance defined as COM of residues K170, Q234, and Y236 and has an RMSD < 1.5 Å to the previous position within the trajectory (see Supplementary results) (15). Density maps of C10 around t-PlaF show that C10 tends to gather in T3 **(Figure 6A)**, as also supported by the finding that T3 is involved in 8.61% of the total binding events of C10 to t-PlaF (**Figure S12**). In most events, the acyl chain of C10 enters T3 first (**Figure 6B**), which is compatible with the mostly hydrophobic interior of T3 (15). T3 has previously been suggested to play a role in the catalytic cycle by relocating substrates during catalysis (15). Thus, the identified C10-PlaF interactions in T3 may influence enzyme activity. Furthermore, it has been hypothesized that FFA released from the GPL may diffuse to the periplasmic space *via* T3 (15), therefore it is feasible that C10 may also enter PlaF *via* the same path.

**Figure 6.**
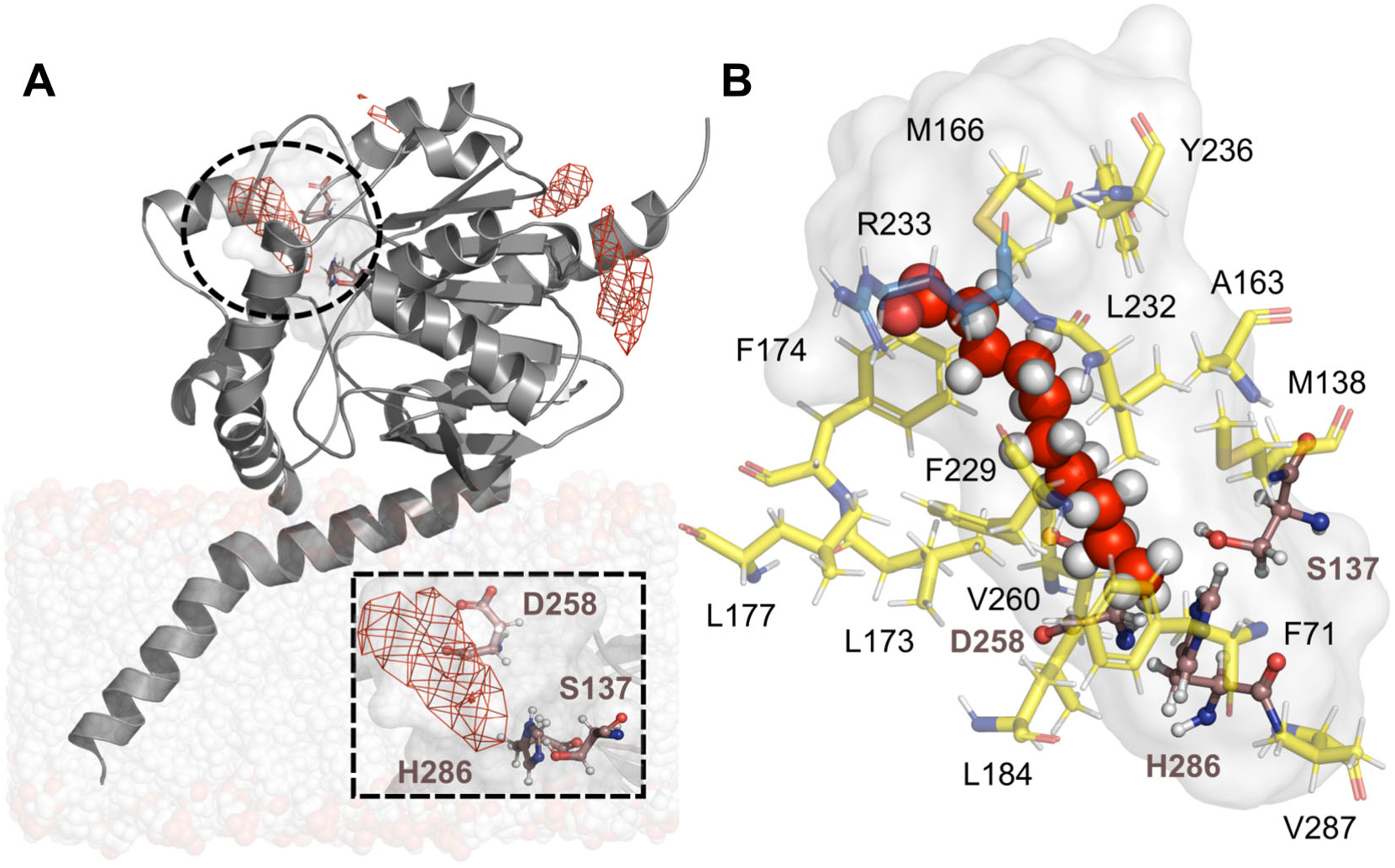
Free-ligand diffusion MD simulations reveal access of C10 molecules to T3. **(A)** 3D density grids (red mash) showing the probability density of C10 around t-PlaF. All C10 were considered in the 3D density grid calculations, and the contour level is set as one standard deviation above the mean value (1σ). T3 computed using CAVER 3.0 (36) is depicted as a light grey surface. The region (black dashed circle) around the catalytic triad (S137, D258, and H286, indicated as brown sticks) is shown in the blow-up (black dashed rectangle). **(B)** A representative binding mode of C10 located within T3 from the most populated cluster that comprises 51.8 ± 0.3% of all bound C10 configurations. T3 was calculated using CAVER 3.0 (36) and is presented as a light grey surface. Carbon and hydrogen atoms of the C10 are depicted as red and grey spheres, respectively. The catalytic triad is illustrated as brown sticks, and the T3 residues as yellow sticks, except for R233 represented as blue sticks.

### 2.7 Tail-first Access of C10 into T3 is Energetically Favorable

As a prerequisite to computing the energetics of binding of C10 to T3 of PlaF, we validated if C10 access *via* its tail is favorable. We applied steered MD (sMD) simulations (37) to pull C10 from the bound state obtained during fldMD simulations to the unbound state in which C10 is located outside T3 in an aqueous milieu. We did not use a trajectory from the fldMD simulations as the C10 tail did not insert deeply enough into T3 to reach the catalytic site. The transition pathway will be later used to define reference points for US simulations to compute a PMF of C10 egress. A bound C10 of replica 4 of the fldMD simulations was chosen as a representative pose (**Table S5** (t-PlaF(III): CAP-1127)). The terminal carbon atom of C10 was considered for pulling from a distance of 3.4 Å to the catalytic oxygen of S137 to 13.1 Å, where it is in the solvent. The C10 was pulled through four consecutive virtual points **(Figure 7A, B)** chosen such that the entire egress pathway was covered as described before (15). The distance between the pulled C-terminal carbon atom and the virtual point was used as a reaction coordinate. The pulling was repeated 50 times for each step, and the work done was computed as a function of the reaction coordinate, as done in previous work (15) **(Figure 7A)**. By applying Jarzynski’s relation (eq. 1) (38), the work was related to the free energy difference between the respective two states along the transition pathway.

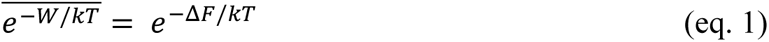

**Figure 7.**
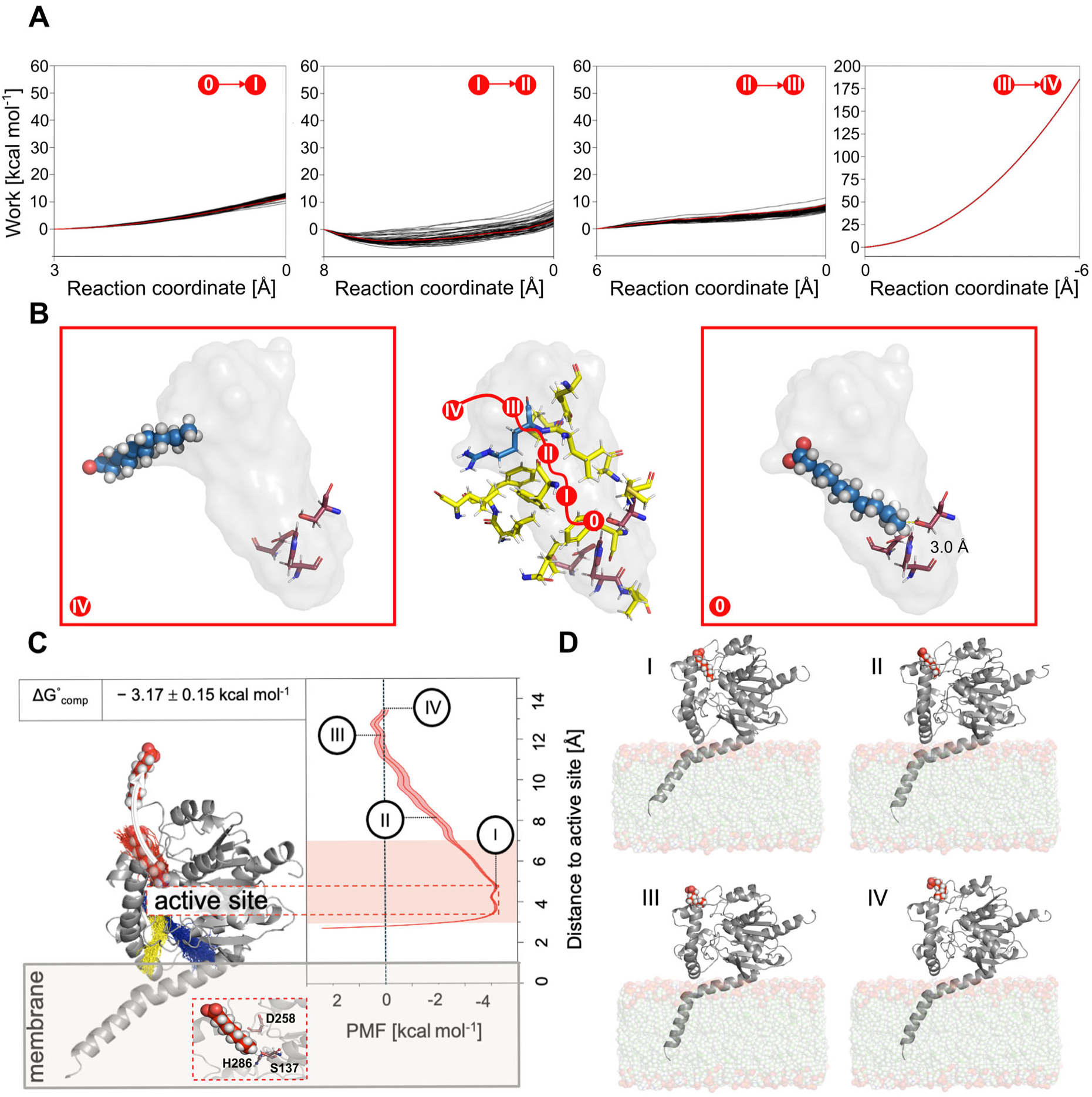
Binding free energy of C10 to T3 as determined by sMD and umbrella sampling. Work distributions (black lines) were obtained from 50 replicas of sMD simulations. **A)** C10 is pulled out of T3 via its tail. C10 is first pulled from the bound position (0) to point I. A replica closest to Jarzynski’s average (red line) was considered as the starting point for the next pulling, I ◊ II. This pulling continues until IV, after which C10 is completely pulled out of T3. The reaction coordinate denotes the distance to the target point. **B)** The starting point and the final point of the pulling are illustrated. (0) is the initial bound position with a distance of 3.0 Å between the catalytic Ser oxygen and the terminal methyl group of the fatty acid. (IV) is the final position, with the terminal methyl group of the fatty acid being pulled out of the tunnel. The catalytic triad residues are represented with purple sticks. The grey T3 tunnel surface was computed with CAVER 3.0 (36). PMF profile of C10 egress through T3 in the t-PlaF configuration **(C)** and corresponding states **(D)**. **(C)** The entrance of T3 (red) is located > 35 Å above the membrane, pointing into the aqueous milieu; T1 and T2 (blue and yellow, respectively) point to the membrane. C10 is pulled from the bound state through T3 into the water phase mimicking the periplasmic space in the cell. The catalytic site is marked with a dotted red box. The red-shaded area around the curve represents the SEM. The red-shaded box corresponds to the integration limits used to calculate *K*_eq_ (eq. 2) to determine Δ*G*^°^_comp_. **(D)** Corresponding states during the C10 egress. State I: Starting position of C10 (in the bound state). State II: C10 tail reaches half of the tunnel. State III: C10 tail is close to the tunnel interface with the periplasmic space. State IV: C10 reaches the end of the T3. The C10 molecule is surrounded by water.

The sMD trajectory whose work-*versus*-reaction coordinate profile is closest to the Jarzynski average was identified as the most favorable transition pathway (15) **(Figure 7B).** Its endpoint provided the starting point for the sMD simulations in the next part of the transition pathway. As a result, the transition pathway is close to the lowest-free energy pathway of C10 egress from the catalytic site to the aqueous milieu. Overall, this approach is similar to sampling unbinding trajectories of ligands from proteins before applying Jarzynski’s relation (39–41) but uses piecewise sMD simulations along the pathway to account for the curvilinear tunnel. A total of 1.15 μs of sMD simulation time was used.

PMFs were computed from US simulations along the sMD-determined transition pathway (24) and post-processed with WHAM (42, 43) to evaluate the energetics of C10 egress. As a reaction coordinate, the distance between the terminal carbon atom of C10 to the side-chain oxygen of the catalytic S137 was used. In total, 29 windows were sampled with 260 ns of sampling time each, of which the first 100 ns were discarded as equilibration. The kernel density showed a median overlap of 33.6 ± 3.1% between contiguous windows, well suited for PMF calculations (26). The PMF of C10 egress is converged and precise **(Figure S13)**. It reveals the presence of an energetic global minimum associated with the bound state I which is ∼4 kcal mol^-1^ more favorable than the partially bound states II and III as well as the unbound state IV (**Figure 7C, D**). The computed binding free energy (Δ*G^°^_comp_*, (25)) of -3.17 ± 0.15 kcal mol^-1^ indicates that tail-first access of C10 to T3 is energetically favorable. These results further strengthen the suggestion that the entry of C10 from the aqueous milieu to T3 and its binding close to S137 could prevent a substrate (GPL or LGPL) from reaching the catalytic site, thereby competitively inhibiting PlaF activity (15).

### 2.8 Tryptophan Substitutions in T3 are suggested to Interfere with C10 Access

To validate the prediction that T3 is the preferred pathway for C10 entering PlaF from the water phase, we used the previously characterized Trp substitutions F229W and L177W within T3 that caused tunnel constriction **(Table 1)** without affecting protein thermostability (15). This strategy had previously been employed to block tunnels of dehalogenases (44) and PlaF (15) with respect to substrate access. T3 could be identified at a similar occurrence rate **(Table 1)** in PlaF_WT_ and both PlaF variants, therefore these Trp variants are suitable for experimentally studying the binding of C10 from the aqueous milieu to the T3.

**Table 1.** Effect of single mutations on C10 binding to T3 in t-PlaF WT and two variants identified from fldMD simulations.

| PlaF | Occurrence<sup>a</sup> | Average bottleneck radius<sup>b,c</sup> | C10 binding events<sup>d</sup> | Effective binding free energy<sup>e</sup> |
| --- | --- | --- | --- | --- |
| WT | 27.75 | $2.29 \pm 0.32$ | 8.61 | $-16.47 \pm 0.53$ |
| F229W | 24.78 | $1.67 \pm 0.31$ | 6.90 | $-17.90 \pm 0.55$ |
| L177W | 28.37 | $1.70 \pm 0.34$ | 5.42 | $-21.28 \pm 1.51$ |

<sup>[a]</sup> Frequency of occurrence of T3 in CAVER; in %.
<sup>[b]</sup> Data was calculated with CAVER using a probe radius of 1.2 Å.
<sup>[c]</sup> In Å; the average and SD are given.
<sup>[d]</sup> Frequency that describes the number of T3-bound frames compared to the total number of frames; in %.
<sup>[e]</sup> In kcal mol<sup>-1</sup>; the average and SEM are given.

MD simulations of free ligand diffusion of C10 were performed to observe binding events to T3 in the two PlaF variants, applying the same criteria as described for PlaF_WT_. The results indicated a reduction in the number of C10 binding events to T3 compared to PlaF_WT_ to 6.90% in PlaF_F229W_ and 5.42% in PlaF_L177W_ variant **(Table 1, Figures S12, S14, S15)**. 3D density grids of C10 molecules from fldMD simulation trajectories indicated that the C10 occurrences around the two engineered Trp sites are reduced compared to the PlaF_WT_ but the orientation and the depth by which C10 immerses into T3 are similar among all three proteins (**Figure 8A**).

**Figure 8.**
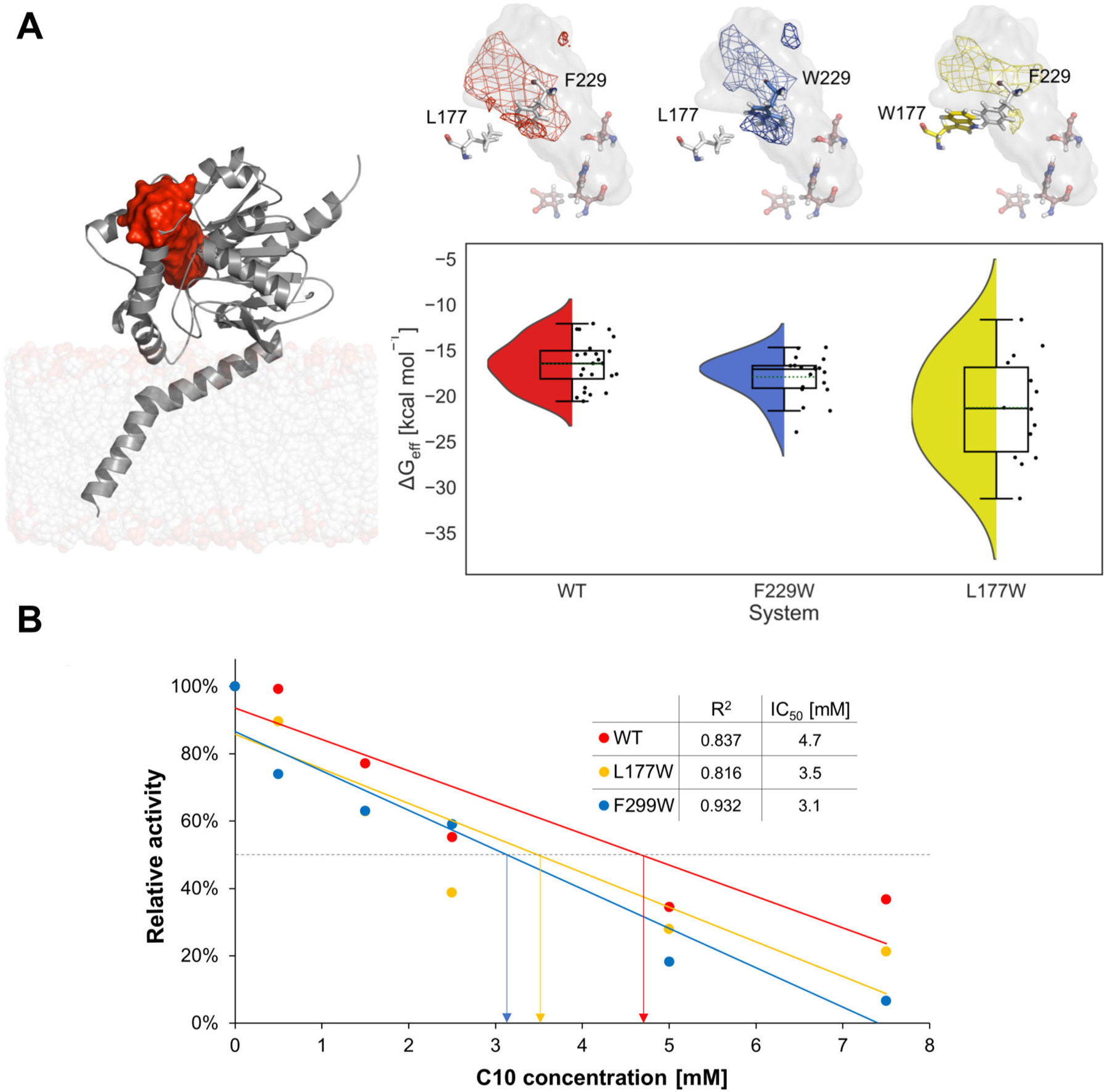
Effect of T3 mutations on FFA binding to PlaF. **A)** Binding of FFA to T3 in the PlaF_WT_ and the Trp variants. Left: The PlaF_WT_ T3 is shown as a red surface in t-PlaF (grey cartoon). Top-right: C10 3D density grids based on fldMD simulations are shown. The contour level is set as 1σ. T3 is depicted as a grey surface and the catalytic triad as brown sticks. The substitution sites are shown in white sticks, except for the included tryptophans, which are represented as blue sticks for PlaF_F229W_ and yellow sticks for PlaF_L177W_. The density grids are colored accordingly, except for the WT, which is shown in red. Bottom-right: The violin plot indicates the distribution of the binding effective energies of FFAs to T3. The inner box of a box plot represents the interquartile range, with the horizontal black line indicating the median. The dotted green line represents the mean, and the vertical black line shows the rest of the distribution, excluding points determined to be outliers when they fall outside 1.5 times the interquartile range. **B)** Inhibitory effect of C10 on PlaF_WT_ and PlaF variants. Linear plots of relative activities (sample without C10 was set to 100%) and C10 at concentrations in a range of 0 – 7.5 mM. Linear correlation coefficients (*R*^2^) and IC_50_ values are indicated. The results represent the mean of initial reaction velocities measured six times using esterase substrate (*p*-NPB) and proteins purified in the presence of DDM.

To further evaluate the effect of the Trp substitutions on C10 interactions within T3, we computed binding effective free energies using MM-PBSA with an implicit membrane model (45, 46). For this, frames from the fldMD simulations with bound C10 were used. The computations are converged, as evidenced by the comparison between the first and the second halves of the trajectories (**Figure S16)**. Across 12 different replicas, we identified 24, 19, and 14 trajectories in which C10 were bound to T3 of PlaF_WT_, PlaF_F229W_, and PlaF_L177W_, respectively. The results revealed that C10 molecules bind to T3 more favorably in the Trp variants (Δ*G*_eff_ (PlaF_F229W_) = -17.90 ± 0.55 kcal mol^-1^; Δ*G*_eff_ (PlaF_L177W_) = -21.28 ± 1.51 kcal mol^-1^) than the WT (Δ*G*_eff_ (PlaF_WT_) = -16.47 ± 0.53 kcal mol^-1^) **(Table 1**, **Figure 8A)**. Such a more prevalent bound state may explain a reduced number of binding events of C10 to T3 of Trp variants (**Table 1**). The more favorable binding in the variants likely arises from better interactions of the lipophilic tail of C10 with the bulky sidechain of Trp than the WT residues.

To experimentally investigate the effect of the Trp substitutions in T3 on the inhibition of PlaF by C10, we produced PlaF_WT_, PlaF_F229W_, and PlaF_L177W_ using the *P. aeruginosa* expression system (6), purified proteins in the presence of dodecyl-β-D-maltosid (DDM) using IMAC (15), and reconstituted them in DOPE:DOPG SUVs using a methodology similar to that employed for LL domain variants. Subsequent inhibition studies by varying concentrations of C10 while keeping the protein and substrate concentrations constant revealed half-maximal inhibitory concentrations (IC_50_) for PlaF_WT_ and Trp variants **(Figure 8B)**. Results show reduced IC_50_ values of Trp variants compared to the WT, indicating a stronger inhibitory effect of C10 on Trp variants in comparison to the PlaF_WT_. These experimental results are in agreement with computed binding effective free energies **(Table 1)** and further strengthen the role of T3 in the binding of C10.

## 3. DISCUSSION

In this study, we combined molecular simulations at the atomistic level with biochemical experiments to examine how the interaction of medium-chain FFAs with PlaF regulates PlaF activity. Using *in vivo*, *in vitro*, and computational analyses, we previously established a model (6) according to which homodimerization of catalytically active PlaF monomers leads to PlaF inhibition, a process induced by medium chain (C10 - C14) FFAs. Following monomerization, PlaF reorients in the bilayer by tilting, (6) which is essential for its activation as it allows membrane-bound GPL substrates to reach the active site (15). Here, we further refined the model of PlaF activity regulation by assessing the impact of C10 located (i) in the GPL bilayer or (ii) in the aqueous milieu surrounding the GPL bilayer on the dimer-monomer transition and the tilting as well as on substrate access to the active site.

### i) Effect of FFAs in the GPL bilayer on PlaF

Our MD simulations and PMF computations revealed that C10 in the GPL bilayer exerts a stabilizing effect on the inactive di-PlaF configuration, in accordance with findings from covalent crosslinking experiments (6). This stabilization likely arises from favorable interactions between FFAs localized in the upper bilayer leaflet and predominantly positively charged residues located in PlaF’s LL domain, which is close to the upper leaflet in the di-PlaF configuration. Mutagenesis experiments and DMP-crosslinking studies using purified proteins functionally validated the predictions by showing that single-charged residue-to-alanine variants exhibited a reduced dimerization compared to PlaF_WT_. Notably, higher C10 concentrations were predicted to stabilize di-PlaF more, which led us to hypothesize that multiple interactions between membrane-localized FFAs and residues within the LL domain occur simultaneously. We validated this hypothesis by using double variants carrying combinations of residues identified as FFA interaction sites in our MD simulations. These double variants showed a more strongly reduced PlaF dimerization than single-point variants. Furthermore, our MD simulations and PMF computations indicated that C10 promotes the t-PlaF configuration and facilitates the tilting process. The latter might be attributed to a potential reduction in membrane viscosity caused by the addition of FFAs as membrane components (47). Still, considering physiological conditions in the *P. aeruginosa* WT and that the equilibria for dimer-to-monomer transition and tilting are coupled, between 2.3 and 7.3% of the PlaF molecules are predicted to be in a monomeric, tilted, catalytically active state in *P. aeruginosa* when 20 mol% C10 are present in the upper leaflet. This is more than 10-fold less than in *P. aeruginosa* WT in the absence of C10 (6). The observed impact of FFAs located in the GPL bilayer on the transitions of PlaF likely contributes to the non-competitive/allosteric component of a mixed inhibition kinetics mechanism previously demonstrated for FFA acting on PlaF (6).

### ii) Effect of FFAs in aqueous milieu on PlaF

Medium-chain FFAs were cocrystallized in the active site of PlaF, and they compete with the substrate according to previous inhibition kinetics results (6). To scrutinize if C10 localized in the water phase of a bilayer-bound PlaF can interact with the enzyme such that substrate binding is impacted, we performed fldMD simulations. These simulations revealed that C10 can bind to the active monomeric t-PlaF *via* T3, which was previously identified to connect the active site of PlaF with the surrounding water milieu (15). C10 favors tail-first access and can reach close to the catalytic triad with its tail. In such a configuration, C10 may potentially interfere with substrate binding, which could explain the experimentally determined competitive effect of C10 on PlaF. We next studied by fldMD simulations previously characterized PlaF variants with genetically engineered bulky Trp residues in T3 (PlaF_F229W_ or PlaF_L177W_) (15). The simulations revealed that binding events for C10 occurred less frequently in either single variant, as might have been expected due to the narrowed T3. Interestingly, binding effective free energy computations revealed that C10 binds stronger to T3 in the variants, i.e., the less frequent binding events there result from C10 being bound longer in T3. These predictions are in line with biochemical studies revealing that inhibition of the PlaF variants reconstituted in SUVs by C10 was enhanced compared to PlaF_WT_. Together, our results indicate that T3 is involved in the inhibition of PlaF by C10 in the water phase.

### Possible physiological relevance of FFA-mediated PlaF inhibition

Comparative GC-MS profiling of FFAs secreted by *P. aeruginosa* Δ*plaF* and the wild-type cells revealed that PlaF can release medium-chain FFAs, including C10 and C14, from GPLs as *in vitro* substrates. This result corroborates our suggestion that PlaF exerts multiple physiological functions (48) through GPL degradation (6). We observed that the chain length of the FFAs has a minor impact on the tilting transition (6), indicating that other medium-chain FFAs may influence PlaF similarly as described here for C10. At physiologically relevant concentrations in *P. aeruginosa*, we estimated that PlaF predominantly exists as inactive di-PlaF in the presence of FFAs. These results together with the predicted function of T3 for C10 inhibition strongly suggest that product feedback regulation of PlaF catalytic activity may be important for *in vivo* PlaF-mediated GPL degradation and membrane remodeling.

In conclusion, our study provides detailed mechanistic insights into the impact of medium-chain FFAs on the regulation of PlaF activity *in vitro*. While likely similar mechanisms are active *in vivo*, this needs to be experimentally validated in the future. Disentangling FFA-mediated activity regulation of the integral membrane protein PlaF is challenging as FFAs may bind into the protein, interact with the protein from the bilayer, and change bilayer properties (e.g., viscosity). Our MD simulations and free energy computations provide evidence that the interplay of these mechanisms serves as a regulatory factor governing PlaF function. Studying membrane proteins in near-native conditions within GPL vesicles as done here shall help to disentangle these complex relationships further. Our results should help understand the regulatory role of FFAs present in the periplasm or inner bacterial membrane on PlaF and eventually other single TM helix spanning membrane proteins because FFAs are among the most common regulators of protein function (49). They also open up new perspectives on how to inhibit PlaF, which has been suggested as a promising target for developing new antibiotics against *P. aeruginosa* (6).

## 4 MATERIALS AND METHODS

### 4.1 Gas Chromatography-mass Spectrometric (GC-MS) Analysis of FAs Extracted from *P. aeruginosa P. aeruginosa*

PA01 (wild-type, WT) and the Δ*plaF* mutant were cultivated overnight (37 °C, LB medium, agitation), and cells were separated from the supernatant, followed by chloroform/methanol extraction of FFAs, derivatization of FFAs with *N*-methyl-N-(trimethylsilyl) trifluoroacetamide, and quantification by GC-MS as described in ref. (50).

### 4.2 Preparation of Starting Structures

The crystal structure of the PlaF dimer is available in the Protein Data Bank (PDB ID 6I8W) (51). The last five residues of the C-terminus of each monomer missing in the structure were added using MODELLER (52), and all small molecule ligands were removed. The dimer was oriented in the membrane using the PPM server (di-PlaF) (22). From that, the “split” s-PlaF_A_ configuration of chain A was generated by removing chain B from the dimer orientation. Additionally, chain A was oriented using the PPM server, resulting in the tilted configuration t-PlaF. These three starting configurations, di-PlaF, s-PlaF, and t-PlaF, were embedded into a DOPE:DOPG = 3:1 membrane (53) and solvated using PACKMOL-Memgen (54, 55). The membrane composition resembles that of the native inner membrane of Gram-negative bacteria (53).

Free fatty acids (FFA) of different chain lengths were added as upper-leaflet components in the protonated form (56) in two different concentrations; here, “upper” refers to the leaflet pointing to the periplasmic space. A distance of at least 15 Å between the protein or membrane and the solvent box boundaries was kept. To obtain a neutral system, counter ions were added that replaced solvent molecules (KCl 0.15 M). The generated systems are summarized in **Table S3**. The size of the resulting systems was ∼140,000 atoms for s-PlaF and t-PlaF and ∼185,000 atoms for di-PlaF.

### 4.3 Unbiased Molecular Dynamics Simulations of PlaF Monomers

The GPU particle mesh Ewald implementation from the AMBER21 suite of molecular simulation programs (57) with the ff14SB (58) and Lipid14/17 forcefields (59, 60) for the protein and the membrane lipids, respectively, were used; water molecules and ions were parametrized using the TIP3P model (61) and the Li and Merz 12-6 ions parameters (62, 63). For the monomer configurations (s-PlaF and t-PlaF), 12 independent simulations of 1 μs length were performed (**Table S3**). Covalent bonds to hydrogens were constrained with the SHAKE algorithm (64) in all simulations, allowing the use of a time step of 2 fs. Details of the thermalization of the simulation systems are given below. All unbiased MD simulations showed stable protein structures and membrane phases evidenced by electron density calculations (**Table S3, Figures S2-S5**).

### 4.4 Relaxation, Thermalization, and Production Runs of the PlaF Monomers

An initial minimization step was performed with the CPU code of pmemd (65). Each minimization was organized in four steps of 1000 cycles each, for a total of 4000 cycles of minimization. Afterward, each minimized system was thermalized in one stage from 0 to 300 K over 25 ps using the NVT ensemble and the Langevin thermostat (66), and the density was adapted to 1.0 g cm^-3^ over 975 ps using the NPT ensemble with a semi-isotropic Berendsen barostat (67) with the pressure set to 1 bar. The thermalization and equilibration were performed with the GPU code of pmemd (65). There are three density equilibration steps with a total time of 4 ns. The sum of thermalization, density adaptation, and equilibration takes 5 ns.

For each replica, 1 μs of production run using the GPU code of pmemd was performed in the NPT ensemble at a temperature of 300 K using the Langevin thermostat (66) and a collision frequency of 1 ps^-1^. To avoid noticeable distortions in the simulation box size, semi-isotropic pressure scaling using the Berendsen barostat (67) and a pressure relaxation time of 1 ps was employed by coupling the box size changes along the membrane plane (68).

### 4.5 Analysis of MD Trajectories of the PlaF Monomer

The trajectories were analyzed with CPPTRAJ (33). As in ref. (6), the angle between the membrane normal and the vector between the COM of Cα atoms of residues 21–25 and residues 35–38 was calculated to describe the tilting of monomeric PlaF.

### 4.6 PMF and Free Energy Calculation of Dimer Dissociation

For calculating a configurational free energy profile (PMF) of dimer separation, 71 intermediate states were generated by separating one chain of the dimer along the membrane plane in steps of 0.5 Å, respectively. The generated structures represent the separation process of the PlaF dimer. To sample configurations along the chain separation in a membrane environment, each intermediate state was embedded into a membrane of approximately 157x157 Å by using PACKMOL-Memgen (55) (see chapter 4.2 and **Table S3** for the composition of the systems). Each intermediate state was minimized, thermalized, and equilibrated following the protocol already described (sections 4.2 and 4.3).

Umbrella sampling simulations were performed starting from each equilibrated intermediate state by restraining the initial distance between chains in every window with a harmonic potential, using a force constant of 4 kcal mol^-1^ Å^-2^ (24); the distance between the COM of C_α_ atoms of residues 25-38 of each monomer was used as a reaction coordinate value *r*. Independent MD simulations of 800 ns in length each were started from each intermediate state, resulting in a total simulation time of 56.8 μs. *r* was recorded every 2 ps and post-processed with the Weighted Histogram Analysis Method implementation of A. Grossfield (WHAM 2.0.9)(42), removing the first 640 ns as an equilibration phase of the system. The error was estimated by considering the last 160 ns only and by calculating the standard error of the mean from eight independent free energy profiles determined every 20 ns during this time. The overlap between contiguous windows and the convergence of the PMFs was validated **(Figures S8-9)**. The association free energy was estimated from the obtained PMF following the membrane two-body derivation (28) and our previous work (6, 27) (eqs. S1-4).

### 4.7 PMF and Free Energy Calculation of Monomer Tilting

The initial conformations used in every window for calculating the PMF of the monomer tilting were obtained from the first microsecond of MD simulations of replica 2 of s-PlaF in the Tilt1 system **(Table S3).** Oriented as in the di-PlaF crystal structure, the monomer spontaneously tilted. The distance *d* along the z-axis between the COM of C_α_ atoms of residues 33-37 of the monomer with the membrane center was used to select 40 intermediate tilting configurations as done already with a standard membrane composition (6). The starting conformations were extracted from the representative trajectory, taking the respective snapshots where *d* showed the least absolute deviation to the average value obtained by binning *d* in windows of 0.5 Å width and with an evenly distributed separation of 0.5 Å. *d* was restrained for every configuration by a harmonic potential with a force constant of 4 kcal mol^-1^ Å^-2^, and sampling was performed for 800 ns per window. *d* values were obtained every 2 ps and analyzed as described above. The error was estimated in the same way as for the dimerization. The overlap between contiguous windows and the convergence of the PMF was validated **(Figure S7)**. The free energy of monomer tilting was estimated from the obtained PMF following protocols explained in previous works (6, 25) (eqs. S5-6).

The dissociation and tilting equilibrium constants as well as the proportion of PlaF dimer versus monomer in a live cell of *P. aeruginosa* were calculated in the same way as in ref. (6); see also the Supplementary Information.

### 4.8 Density Maps of Free Fatty Acids Binding to di-PlaF in the Membrane

The structure of the PlaF dimer was embedded in a phospholipid membrane with the composition DOPE:DOPG 3:1 following the protocol described in section 4.1. C10 fatty acids were added as membrane upper leaflet components in the protonated configuration (56). The final composition of the upper leaflet is DOPE:DOPG:C10 3:1:1. Unbiased MD simulations of the system were performed following the protocol used for the monomers (see sections 4.2 and 4.3). From the obtained trajectories, all FFA binding poses were identified in which FFAs have an RMSD < 1.5 Å to the previous frame (see Supplementary results), and are located at most 5 Å away from the protein. These binding poses were clustered using the hierarchical agglomerative (bottom-up) algorithm implemented in cpptraj (33), using the minimum distance *ε* between the clusters as the cluster criterion. Starting from *ε* = 2.0 Å, we gradually increased *ε* in 0.5 Å intervals until the population of the largest cluster remained unchanged, (*ε* = 5.0 Å). We calculated the 3D density maps of the fatty acids considering all atoms using the grid function available in CPPTRAJ, using a grid spacing of 1.5 Å (33). We applied a contour level of 1σ (one standard deviation above the mean value).

### 4.9 fldMD to study C10 FFA Binding from the Water Phase to t-PlaF

t-PlaF was embedded in a phospholipid membrane with the composition DOPE:DOPG = 3:1 following the protocol described in section 4.1. To investigate the binding of FFAs to PlaF, 1 µs long MD simulations of the free diffusion of FFA molecules located in the water phase were performed to investigate their binding to t-PlaF. The FFAs were assumed to be in their deprotonated configuration, resulting in a concentration of ∼30 mM in the corresponding solvation box. The eventual formation of aggregates was checked by visual inspection of the trajectories. 12 replicas of 1 µs simulation length were run for each system. The minimization, thermalization, equilibration, and production schemes are like the ones performed in sections 4.2 and 4.3.

### 4.10 Simulated Extraction of Fatty Acids through T3

MD simulations were performed using the GPU implementation of the AMBER 21 molecular simulation package (55), employing the same protocol as in chapters 4.3 and 4.4. To extract an FFA from T3 into the periplasmic space, we selected a bound FFA (**Table S3** (t-PlaF(III): CAP-1127)) of replica 4 of the fldMD simulations as a representative pose. We pulled it by its terminal carbon atom from the catalytic center to the tunnel’s exit using constant velocity sMD simulations, with a constant velocity of 1 Å ns^-1^ and a force constant of 5 kcal mol^-1^ Å^-2^ (37, 41). Pulling simulations at low velocities have been used with lipids to calculate free energy profiles (15, 69). Using low pulling rates, the lipids have time to adapt to energetically favorable conformations during the extraction process. T3 was divided into consecutive fragments connected to T3 pulling points generated by the COM of specific amino acid residues (**Table S5**). We performed 50 replicas for each pulling simulation to identify the lowest energy pathway. The work was computed as a function of the reaction coordinate. The computed work was further related to the free energy difference between two states of the pulling simulation by applying Jarzynski’s relation (eq. 1) (38). Δ*F* is the free energy difference between two states, which is connected to the work *W* done on the system. *k* is the Boltzmann constant, and *T* is the absolute temperature of the system. The replica closest to Jarzynski’s average (38) was considered to describe the lowest free energy pathway, and provided the starting point for the next pulling stage, as done in ref. (15). This procedure results in faster convergence of PMF profiles subsequently computed along the pathway, decreasing the overall computations needed (39).

### 4.11 Umbrella Sampling Simulations and PMF calculations

To understand the energetic contribution associated with the binding of C10 molecules to T3, PMFs were computed based on umbrella sampling, taking structures from the sMD simulations (see chapter 4.10) as starting points. As a reaction coordinate, the distance of the terminal carbon atom of the FFA to the hydroxyl oxygen of S137 of the active site was used. Consecutive positions of the FFA from the bound state to the periplasmic space, as determined in chapter 4.10, were considered reference points, defining the umbrella windows. To achieve sufficient overlap between the umbrella windows, distances between reference points of 0.5 Å were used. The length of T3 and the size of the FFA can vary. Therefore, for sampling the egress of the bound fatty acid, different numbers of windows were required for the tunnel. The FFA was restrained by harmonic potentials to the reference points, using a force constant of 5 kcal mol^−1^ Å^−2^. To achieve sufficient convergence of the PMF profile **(Figure S14)**, each window was sampled for 260 ns, of which the last 160 ns were used to calculate the PMF. Distance values were recorded every 2 ps and processed with WHAM (42). The PMFs were evaluated for convergence by checking the change in the free energy profile with increasing sampling time in steps of 20 ns **(Figure S14)**.

### 4.12 Absolute Binding Free Energy from the PMF

The absolute binding free energy of fatty acids to PlaF was determined from the computed PMF using an approach modified (15) from Chen and Kuyucak (26). The PMF was integrated along the reaction coordinate (eq. 2) to calculate an association (equilibrium) constant (*K*_eq_).

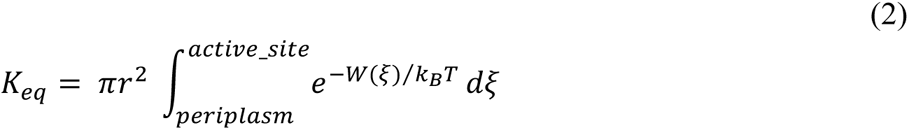

Here, *r* is the maximum bottleneck radius of the respective tunnel, which was determined by CAVER analysis (15, 36), π*r*^2^ is the maximum cross-sectional area of the tunnel, *W*(ξ) is the PMF at a specific value of the reaction coordinate, *k* is the Boltzmann constant, and *T* is the absolute temperature at which the simulations were performed. The integration limits describe the bound state. A reaction coordinate > 7 Å was considered as an unbound state and for this reason not included in the integration.

*K*_eq_ was then transformed to the mole fraction scale (*K*_x_). For (un)binding to the solution, a standard state of 1 M is considered (25) (eq. 3).

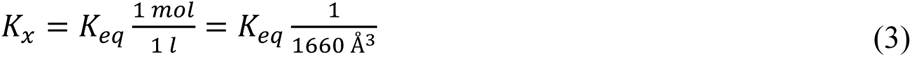

From *K*_x_, the difference in the standard free energy (eq. 4) between the bound and unbound states (Δ*G*_comp_^°^) of a single substrate molecule was calculated.

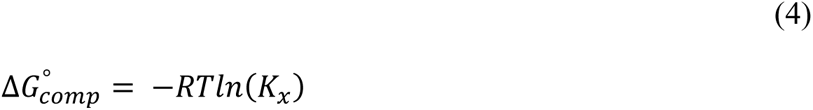

### 4.13 Predicting Mutations to Modify Access of the Free Fatty Acid to T3

To verify the prediction that FFA can bind to T3 and, thereby, inhibit PlaF, we intended to alter the geometry of T3 by introducing small-to-tryptophan substitutions of tunnel-lining residues, following an approach already pursued (15). The selected residues were substituted to tryptophan using FoldX (70), and the stability of the PlaF variants was evaluated in terms of the change in folding free energy (ΔΔ*G*) with respect to the wild-type (71) **(Table S6)**. Single and double amino acid substitutions were performed 10 times for each residue or residue couple, and the results were averaged. If the average ΔΔ*G* > 3 kcal mol^-1^, the substitution is considered destabilizing (72) and was not further pursued. To check if the proposed substitutions will modify the tunnel geometry, the bottleneck radius and the tunnel length of the variant tunnels were calculated using CAVER 3.0 (36) **(Table S6)**. The COM of the catalytic residues S137 and H286 were defined as the starting points of the search, as done in our previous work (15). The probe radius was set to 1.2 Å.

Mutant fldMD simulations and binding free energy calculations of C10 to T3 were performed by following the same protocols described for the wild-type PlaF.

### 4.14 MM-PBSA Calculations of Free Fatty Acid Binding to t-PlaF from the Water Phase

To pinpoint the most likely binding epitopes, we generated 3D density grids to map the location of each fatty acid. Conformations stably bound (as defined in chapter 4.8) were then clustered to extract the most representative binding poses. Subsequently, we conducted MM-PBSA (Molecular Mechanics-Poisson Boltzmann Surface Area) calculations to determine the binding effective energy of FFAs binding to t-PlaF from the water phase. These calculations were performed for both the WT PlaF and the proposed mutants, utilizing the trajectories from fldMD. FFA binding poses were identified as those exhibiting an RMSD < 1.5 Å compared to the previous frame (as described in chapter 2.5). Additionally, these poses were required to be located within 5 Å of the entrance of T3.

To compute the average binding effective energy of each FFA interacting with t-PlaF within the ensemble of C10 frames, we employed MMPBSA.py (46). This was done using dielectric constants of 1.0 for the protein, 80.0 for the solvent, and 15.0 for the membrane (73). A heterogeneous dielectric model was used to represent the membrane; the implicit membrane model using spline fitting (memopt = 3) was employed for these calculations (45).

### 4.15 Cloning, Site-Directed Mutagenesis, Expression, and Purification

Molecular biology methods were performed following previously described procedures (6). For obtaining PlaF variants of the lid-like domain, site-directed mutagenesis of *plaF* (*pa2949*) was carried out using the Quik-Change PCR method, with the Pfu DNA polymerase and the pET_*pa2949* plasmid (74). Similarly, mutants located in the T3 tunnel were generated through site-directed mutagenesis on pBBR1MCS-3_pa2949 (29). Successful site-directed mutagenesis was confirmed by DNA sequencing. For the production of PlaF and the lid-like domain variants, *E. coli* BL21(DE3) cells transformed with the respective expression vectors were grown overnight at 37°C in lysogeny broth (LB) medium (75) supplemented with ampicillin (100 µg/ml) (74). These cultures were used to inoculate an expression culture in “autoinduction” medium (terrific broth medium containing 0.04% lactose (w/v) and 0.2% glucose (v/v)) supplemented with ampicillin (100 µg/ml) to an initial OD_600nm_ = 0.01. The cultures were then grown for 24 h at 37 °C and harvested by centrifugation at 6750×*g* and 4 °C for 15 min.

For the production of PlaF and T3 tunnel variants, transformed *P. aeruginosa* PA01 cells were grown overnight at 37°C in LB medium supplemented with tetracycline (100 µg/ml) (6). These cultures were used to inoculate an expression culture in LB medium supplemented with tetracycline (100 µg/ml) to an initial OD_600nm_ = 0.05. The cultures were grown at 37 °C until reaching OD = 2, and then harvested by centrifugation at 6750×*g* and 4 °C for 15 min.

The total membrane fraction isolated by ultracentrifugation was solubilized with Triton X-100 (6) and proteins were purified using Ni-NTA IMAC and buffers supplemented with 20 mM OG for lid-like protein variants and 0.25 mM DDM for T3 tunnel protein variants. For biochemical analysis, the proteins were transferred to Tris-HCl buffer (100 mM, pH 8) supplemented with the respective detergents. The PlaF protein and its variants were analyzed by sodium dodecyl sulfate-polyacrylamide gel electrophoresis (SDS-PAGE) under denaturation conditions on a 12 % (v/v) gel (76).

### 4.16 Reconstitution of Proteins into GPL SUVs

Small unilamellar vesicles (SUV) were constructed and used for reconstitution according to a modified protocol (77). The GPLs used, 1,2-dioleoyl-*sn*-glycero-3-phosphoglycerol (DOPG) and 1,2-dioleoyl-*sn*-glycero-3-phosphoethanolamine (DOPE), were dissolved in chloroform (Avanti Polar Lipids, Alabaster, USA).

To produce 2.6 µmol SUVs for protein reconstitution and 64.9 µmol SUVs for fatty acid reconstitution, DOPE and DOPG were mixed in a 1:1 ratio in a glass reaction vessel. The GPLs were dried under a gentle stream of nitrogen and by centrifugation under a vacuum for 20 min. Subsequently, HEPES buffer (20 mM, pH 8, 100 mM NaCl) was added to the dried GPLs and incubated for 10 min at room temperature. The GPLs were then vortexed and sonicated for 2 min. OG and DDM were added to the SUVs at a detergent:SUV ratio of 2:1 or 1:1 (mol/mol) to destabilize the SUVs, respectively. After destabilization of SUVs, samples were incubated with rotation for 1 h to ensure complete equilibration of the detergent with the lipids.

FAs were mixed with the respective detergent-destabilized SUVs to achieve a GPL-to-FA ratio of 5:1 (mol/mol). Proteins were mixed with the respective detergent-destabilized SUVs, with or without C10 to achieve a GPL-to-protein mass ratio of 20:1. Final protein concentration were 1000 nM to achieve dimerization. To remove detergent, approximately 20 activated polystyrene Bio-Beads were added to the reconstitution solution and incubated for 1 h with rotation at room temperature.

### 4.17 Cross-Linking Assays

*In vitro* cross-linking using the bifunctional cross-linking reagent dimethyl pimelimidate (DMP) was performed as previously described (6). Briefly, 30 µl of proteins reconstituted into SUVs were incubated with 1.8 µl of decanoic acid (332 mM) and 18 µl of freshly prepared DMP in phosphate buffer saline (pH 7.4, 150 mM) for 2 h at room temperature to ensure partitioning of C10 to the bilayer. The cross-linking reaction was terminated with 15 µl of stop solution (50 mM Tris-HCl, 1 M glycine, 150 mM NaCl, pH 8.3).

### 4.18 Enzyme Activity Assay

The esterase activities of SUV-reconstituted PlaF and T3 variants were determined at 37 °C, using *p*-nitrophenyl butyrate (*p*-NPB) as a substrate, following the protocol previously described (78). The protein (5 µl) was mixed with 93 µl of freshly prepared 1 mM *p*-NPB solution and 2 µl of C10 dissolved in DMSO. Final protein concentration were 130.22 nM, 280.52 nM, 77.22 nM for PlaF_WT_, PlaF_L177W_, PlaF_F299W_, respectively, and final FA concentrations were 0.5, 1.5, 2.5, 5, and 7.5 mM. Activities were determined by measuring the release of *p*-nitrophenol spectrophotometrically during 1 h at 410 nm. Inhibition was assessed by calculating the relative activities of the inhibited samples in comparison to the activity of the sample without C10, which was set to 100%. IC_50_ values were estimated from linear plots as done in ref. (79).

### 4.19 Thermal Stability

PlaF and variants loaded into the measuring capillaries (Prometheus NT.Plex nanoDSF Grade Standard Capillary Chips) were heated from 20 to 90 °C (1 °C/min, heating rate), and the intrinsic protein fluorescence was recorded at 330 and 350 nm using the Prometheus NT.Plex nanoDSF device (NanoTemper, Munich, Germany).

### 4.20 SDS-PAGE and Western Blotting

Proteins were analyzed by sodium dodecyl sulphate-polyacrylamide gel electrophoresis (SDS-PAGE) under denaturation conditions on 12% (w/v) gels as described previously (74). The proteins were transferred from SDS-PAGE gel to the polyvinylidene difluoride membranes by western blotting and were detected using anti-His(C-term)-HRP antibodies (Invitrogen) as described previously (80).

### 4.21 Detergent quantification

We quantified detergent following established procedures (81). To do so, 25 µl of each protein sample, 50 µl 5% phenol, and 125 µl 96% sulfuric acid were pipetted in a 1.5 ml Eppendorf tube. Samples were vortexed and incubated at 90°C for 5 minutes. After the samples were cooled down at room temperature, 150 µl were transferred to a 96-well microtiter plate and absorbance was measured at 490 nm. The calibration curve was prepared with respective detergent in the same manner as the samples.

## 5 ACKNOWLEDGEMENTS

This study was funded by the Deutsche Forschungsgemeinschaft (DFG, German Research Foundation) project no. 267205415 / CRC1208 grant to F.K. and K.-E.J. (subproject A02) and H.G. (subproject A03). We thank Muttalip Caliskan (IMET, HHU Düsseldorf) for help with fatty acid quantification. We are grateful for computational support by the “Zentrum für Informations und Medientechnologie” at the Heinrich-Heine-Universität Düsseldorf and the computing time provided by the John von Neumann Institute for Computing (NIC) to H.G. on the supercomputer JUWELS at Jülich Supercomputing Centre (JSC) (user IDs: HDD18; plaf).

## 6 AUTHOR CONTRIBUTIONS

R.G. computational investigation, analysis, visualization; M.M., B.T. experimental investigation, analysis, visualization; S.S. supervision, analysis; K.-E.J., F.K., H.G. conceptualization, supervision, analysis, funding, resources, project management. The manuscript was written with the contributions of all authors. All authors have given approval for the final version of the manuscript.

