## Supporting Information for "Molecular Mechanisms Underlying Medium-Chain Free Fatty Acid-regulated Activity of the Phospholipase PlaF from *Pseudomonas aeruginosa*"

### Supplementary results

*Potential of mean force and free energy computations of dimer association*

The PMF of dimer association is integrated along the reaction coordinate to calculate an association constant ($K_a$), which is subsequently transformed into the mole fraction scale ($K_x$). This transformation considers the number of lipids $N_L$ per surface area $A$, as described in refs. (1, 2). To calculate the difference in free energy between dimer and monomers ($\Delta G$), we utilize the following equations (eq. S1-3):

$$K_a = \frac{||\Omega||}{(2\pi)^2} \int_0^D r e^{\frac{-w(r)}{k_B T}} dr \quad \text{(eq. S1)}$$

$$K_x = K_a \frac{N_L}{A} \quad \text{(eq. S2)}$$

$$\Delta G = -RT \ln(K_X) \quad \text{(eq. S3)}$$

$r$ is the value of the reaction coordinate, $w(r)$ is the PMF at value $r$, $D$ is the maximum distance at which the protein is still considered a dimer, $k_B$ is the Boltzmann constant, and $T$ is the absolute temperature at which the simulations were performed. The factor $||\Omega||/(2\pi)^2$ takes into account the restriction of the configurational space upon dimer formation, in terms of the sampled angle between the two chains in the dimeric state ($||\Omega||$, eq. S4) and the accessible space for the monomers, $(2\pi)^2$.

$$||\Omega|| = [\max(\theta_a) - \min(\theta_a)] * [\max(\theta_b) - \min(\theta_b)] \quad \text{(eq. S4)}$$

In eq. S4, $\theta_a$ is defined as the angle formed between the vectors connecting the COM of chain b with the COM of chain a, with the COM computed from residues 25-38 of the latter chain. $\theta_b$ is defined analogously starting from the COM of chain a. A value for $||\Omega||$ of 0.146 computed from eq. S4 indicates the fraction of accessible space that the PlaF monomers have in the dimeric state compared to when both chains rotate independently.

*Potential of mean force and free energy computations of monomer tilting*

For calculating the free energy difference between the obtained basins, the PMF of monomer tilting was integrated using eq. S5 and is used to calculate the difference in free energy between tilted and straight monomers ($\Delta G_{tilting}$) (eq. S6), according to ref. (3):

$$K_{tilting} = \frac{\int_{B1} e^{-\frac{w(d)}{k_B T}} dr}{\int_{B2} e^{-\frac{w(d)}{k_B T}} dr} \qquad \text{(eq. S5)}$$

$$\Delta G_{tilting} = -RT \ln K_{tilting} \qquad \text{(eq. S6)}$$

$d$ is the value of the reaction coordinate, $w(d)$ is the value of the PMF at that distance, and $B_1$ and $B_2$ represent the basins for the tilted and split configurations of PlaF, respectively. The integration limits $B_1$ and $B_2$ included the flat regions with the minimum slope that indicate the split and the tilted configurations of PlaF, respectively **(Figure S5)**.

*Equilibrium of the PlaF dimer versus monomer under in vivo conditions*

The equilibrium between the PlaF dimer and monomer within the cellular membrane is the result of the interplay between the following equilibria (1):

$$2M \overset{K_a}{\leftrightharpoons} D \quad K_a = \frac{[D]}{[M]^2} \qquad \text{(eq. S7)}$$

$$M \overset{K_{tilting}}{\leftrightharpoons} M_{tilted} \quad K_{tilting} = \frac{[M_{tilted}]}{[M]} \qquad \text{(eq. S8)}$$

This leads to the following relationship:

$$D \overset{K_a\, K_{tilting}^{-2}}{\rightleftharpoons} 2M_{tilted} \qquad \text{(eq. S9)}$$

*D* represents the PlaF dimer, *M* denotes the ‘split’ monomer, and $M_{\text{tilted}}$ represents the tilted monomer. $K_{\text{a}}$ (eq. S1) and $K_{\text{tilting}}$ (eq. S5) are the equilibrium constants for dimer association and monomer tilting in the presence of free fatty acids, obtained from PMF calculations.

We consider the concentrations of the PlaF monomers under overexpressing and non-overexpressing conditions as in our previous study (1). Assuming an approximate lipid area of 63 $Å^2$ per leaflet (or 31.5 $Å^2$ in a bilayer) for a DOPE:DOPG:C10//DOPE:DOPG=3:1:1//3:1 membrane at 300 K, the total area concentration of PlaF molecules is:

$$T = 2[D] + [M] = [\,1.67\,x\,10^{-8}\,, 1.67\,x\,10^{-7}]\,\frac{Plaf}{Å^2} \qquad \text{(eq. S10)}$$

Expressing the association constant in terms of the monomer concentration using eq. S7 yields

$$K_a = \frac{\frac{T-[M]}{2}}{[M]^2} \quad \Leftrightarrow \quad 2K_a[M]^2 + [M] - T = 0 \qquad \text{(eq. S11),}$$

Solving the quadratic equation results in:

$$[M] = \frac{-1+\sqrt{1+8K_aT}}{4K_a} = [\,7.05\,x\,10^{-12}, 2.23\,x\,10^{-11}]\,\frac{PlaF}{Å^2} \qquad \text{(eq. S12)}$$

And:

$$[D] = \frac{T-[M]}{2} = [8.35\,x\,10^{-9}, 8.35\,x\,10^{-8}]\,\frac{PlaF_{dimer}}{Å^2} \qquad \text{(eq. S13)}$$

These results indicate that in live cells, in the presence of FFAs, the fraction of PlaF in the dimeric (monomeric) state is between 99.96 % and 99.99 % (0.04 % and 0.01 %), where the PlaF monomer is in the "split" configuration. In the absence of FFAs, the fraction of PlaF in the dimeric (monomeric) state was between 35 and 72% (65 and 28%) (1).

Since the tilting of the PlaF monomer is energetically favorable compared to the "split" configuration and reduces the concentration of "split" PlaF monomers, the concentration of dimeric PlaF will further decrease. To quantitatively consider the effect of tilting in the presence of FFAs, we express the overall equilibrium constant as:

$$K = K_a \, K_{tilting}^{-2} = \frac{[D]}{[M_{tilted}]^2} \qquad \text{(eq. S14)}$$

Where:

$K_{tilting} = \frac{[M_{tilted}]}{[M]} = 173.27$, equivalent to $\Delta G_{\text{tilting}} = -3.07 \frac{\text{kcal}}{\text{mol}}$, calculated according to eq. S5.

Following the same procedure as before (1), we obtain:

$$[M_{tilted}] = [3.86 \times 10^{-9}, 1.22 \times 10^{-9}] \frac{\text{PlaF}}{\text{Å}^2} \qquad \text{(eq. S15)}$$

$$[D] = [8.36 \times 10^{-8}, 8.35 \times 10^{-9}] \frac{\text{PlaF dimer}}{\text{Å}^2}, \qquad \text{(eq. S16)}$$

These results demonstrate that in live cells, in the presence of FFAs at a ratio of DOPE:DOPG:FFA 3:1:1 in the upper membrane leaflet, the fraction of PlaF in the tilted monomeric (dimeric) state ranges from 2.3 to 7.3 % (97.7 to 92.7 %).

*Determination of the binding mode models for C10 FFAs.*

During visual inspection of the MD trajectories, we observed multiple instances of C10 FFA molecules binding to and unbinding from t-PlaF. To identify bound fatty acid poses to T3, we employed a distance cut-off of 5 Å from the tunnel entrance, defined by the COM of residues K170, Q234, Y236. Subsequently, we determined stably bound fatty acid poses by calculating the RMSD of each fatty acid molecule concerning its previous configuration after superimposing the t-PlaF structure, using cpptraj

(4). This allowed us to use RMSD as a measure of the spatial displacement of a fatty acid molecule between two consecutive snapshots. We considered C10 poses with an RMSD of less than 1.5 Å between consecutive frames as stably bound.

Beforehand, we computed the mean square displacement $<\Delta^2 r(t)>$ for C10 to estimate the spatial displacement for free diffusive motion, according to eq. S16 (5):

$$<\Delta^2 r(t)> \sim 6\, Dt \qquad \text{(eq. S16)}$$

$D$ is the diffusion coefficient C10 in $H_2O$ solution at 20°C: $D = 2.33\ ^{*}\ 10^{-10}\ m^2\ s^{-1}$; this value was calculated with the Cytiva calculator at cytivalifesciences.com (accessed on 09.12.2022). $t$ represents the time interval of saving atomic positions into a trajectory file ($t = 200$ ps). Note that eq. S16 is valid only if $t >> t_c$ ($t_c$: molecular correlation time). Solving eq. S16 yields for C10 $\sqrt{< \Delta 2r(t) >}$ = 5.3 Å. RMSD values smaller than $\sqrt{< \Delta 2r(t) >}$ result from restrictions on the diffusive motion of the fatty acid molecule during the MD simulations, primarily due to complex formation with t-PlaF.

The stably bound poses were further analyzed using pose clustering. We employed the hierarchical agglomerative (bottom-up) algorithm as implemented in cpptraj (4). Initially, we set the minimum distance $\varepsilon$ between the clusters to $\varepsilon = 2.0$ Å and gradually increased $\varepsilon$ in 0.5 Å intervals. We continued this process until the population of the largest cluster remained unchanged, which occurred at $\varepsilon = 5.0$ Å.

### Supplementary tables

**TABLE S1. Medium chain GPLs identified in *P. aeruginosa* WT and Δ*plaF***

| GPL | *P. aeruginosa* PAO1[a] | *P. aeruginosa* Δ*plaF*[a] | *p*-value[b] |
|---|---|---|---|
| PE 15:0 | 0.0117 ± 0.0063 | 0.0081 ± 0.0039 | 0.37 |
| PG 15:0 | 0.0034 ± 0.0016 | 0.0030 ± 0.0012 | 0.71 |
| PE 17:0 | 0.0026 ± 0.0009 | 0.0022 ± 0.0026 | 0.78 |
| PG 13:0 | 0.0014 ± 0.0006 | 0.0013 ± 0.0005 | 0.80 |
| PG 17:0 | 0.0013 ± 0.0001 | 0.0009 ± 0.0010 | 0.50 |
| PG 16:0 | 0.0011 ± 0.0005 | 0.0006 ± 0.0002 | 0.14 |
| PG 23:1 | 0.0010 ± 0.0007 | 0.0010 ± 0.0005 | 0.97 |
| PG 24:2 | 0.0010 ± 0.0003 | 0.0011 ± 0.0007 | 0.89 |
| PG 21:1 | 0.0010 ± 0.0005 | 0.0008 ± 0.0004 | 0.67 |
| PE 19:0 | 0.0009 ± 0.0002 | 0.0011 ± 0.0005 | 0.51 |
| PE 8:0 | 0.0009 ± 0.0004 | 0.0009 ± 0.0008 | 0.87 |
| PG 19:0 | 0.0008 ± 0.0006 | 0.0003 ± 0.0003 | 0.26 |
| PE 18:0 | 0.0007 ± 0.0006 | 0.0010 ± 0.0006 | 0.57 |
| PE 12:0 | 0.0007 ± 0.0008 | 0.0009 ± 0.0011 | 0.74 |
| PE 14:0 | 0.0007 ± 0.0005 | 0.0008 ± 0.0006 | 0.80 |
| PG 12:0 | 0.0007 ± 0.0007 | 0.0004 ± 0.0005 | 0.49 |
| PE 16:0 | 0.0006 ± 0.0005 | 0.0011 ± 0.0006 | 0.30 |
| PE 24:3 | 0.0006 ± 0.0006 | 0.0006 ± 0.0003 | 0.99 |
| PG 22:1 | 0.0006 ± 0.0005 | 0.0012 ± 0.0008 | 0.25 |
| PE 23:1 | 0.0006 ± 0.0004 | 0.0005 ± 0.0000 | 0.71 |
| PE 24:1 | 0.0006 ± 0.0003 | 0.0007 ± 0.0004 | 0.68 |
| PG 9:0 | 0.0006 ± 0.0003 | 0.0009 ± 0.0004 | 0.21 |
| PG 18:0 | 0.0006 ± 0.0003 | 0.0006 ± 0.0005 | 0.87 |
| PE 13:0 | 0.0006 ± 0.0003 | 0.0009 ± 0.0002 | 0.15 |
| PG 19:1 | 0.0006 ± 0.0002 | 0.0005 ± 0.0005 | 0.89 |
| PE 24:2 | 0.0006 ± 0.0008 | 0.0004 ± 0.0003 | 0.81 |
| PG 23:0 | 0.0005 ± 0.0003 | 0.0005 ± 0.0004 | 0.96 |
| PE 21:0 | 0.0005 ± 0.0003 | 0.0005 ± 0.0004 | 0.94 |
| PE 22:2 | 0.0005 ± 0.0003 | 0.0013 ± 0.0018 | 0.46 |
| PE 20:1 | 0.0005 ± 0.0004 | 0.0008 ± 0.0007 | 0.44 |
| PG 24:0 | 0.0005 ± 0.0002 | 0.0006 ± 0.0002 | 0.63 |
| PE 23:2 | 0.0005 ± 0.0002 | 0.0012 ± 0.0006 | 0.09 |
| PG 14:0 | 0.0005 ± 0.0004 | 0.0007 ± 0.0004 | 0.48 |
| PG 21:0 | 0.0005 ± 0.0003 | 0.0004 ± 0.0002 | 0.77 |
| PG 24:1 | 0.0005 ± 0.0002 | 0.0005 ± 0.0004 | 0.98 |
| PG 20:0 | 0.0005 ± 0.0002 | 0.0004 ± 0.0005 | 0.96 |
| PG 20:1 | 0.0004 ± 0.0004 | 0.0009 ± 0.0005 | 0.20 |
| PE 21:2 | 0.0004 ± 0.0003 | 0.0009 ± 0.0006 | 0.24 |

| | | | |
|---|---|---|---|
| PG 23:2 | 0.0004 ± 0.0001 | 0.0001 ± 0.0001 | *0.01* |
| PE 19:1 | 0.0004 ± 0.0004 | 0.0005 ± 0.0003 | 0.72 |
| PG 10:0 | 0.0004 ± 0.0003 | 0.0004 ± 0.0003 | 0.80 |
| PE 10:0 | 0.0003 ± 0.0002 | 0.0002 ± 0.0001 | 0.29 |
| PG 22:0 | 0.0003 ± 0.0002 | 0.0004 ± 0.0003 | 0.71 |
| PE 23:0 | 0.0003 ± 0.0002 | 0.0006 ± 0.0006 | 0.35 |
| PE 9:0 | 0.0003 ± 0.0001 | 0.0005 ± 0.0004 | 0.46 |
| PE 20:0 | 0.0003 ± 0.0001 | 0.0004 ± 0.0001 | 0.46 |
| PE 24:0 | 0.0003 ± 0.0001 | 0.0005 ± 0.0001 | 0.11 |
| PG 22:2 | 0.0003 ± 0.0002 | 0.0005 ± 0.0004 | 0.49 |
| PG 8:0 | 0.0003 ± 0.0001 | 0.0004 ± 0.0002 | 0.55 |
| PG 21:2 | 0.0003 ± 0.0001 | 0.0004 ± 0.0003 | 0.33 |
| PE 11:0 | 0.0002 ± 0.0001 | 0.0003 ± 0.0003 | 0.63 |
| PE 21:1 | 0.0002 ± 0.0001 | 0.0005 ± 0.0006 | 0.38 |
| PG 24:3 | 0.0002 ± 0.0001 | 0.0005 ± 0.0001 | *0.03* |
| PE 22:0 | 0.0002 ± 0.0001 | 0.0007 ± 0.0005 | 0.17 |
| PE 22:1 | 0.0001 ± 0.0001 | 0.0006 ± 0.0002 | *0.02* |
| PG 11:0 | 0.0001 ± 0.0001 | 0.0002 ± 0.0002 | 0.55 |

[a] In nmol mg$^{-1}$ (GPL) / OD580 nm ± SD.

[b] *t*-test (WT vs. Δ*plaF*); values in italics indicate $p < 0.05$.

**TABLE S2. Total PL amount and amount of medium-chain length PLs in *P. aeruginosa* WT and Δ*plaF*.**

| | *P. aeruginosa* WT [nmol/mg(GPL)/OD580nm] ± SD | *P. aeruginosa* Δ*plaF* [nmol/mg(GPL)/OD580nm] ± SD |
|---|---|---|
| Medium chain GPLs* | 0.39 ± 0.19 | 0.42 ± 0.25 |
| All GPLs** | 3.10 ± 0.78 | 2.01 ± 0.70 |
| | 12.6% medium-chain | 20.8% medium-chain |

**TABLE S3. Membrane systems generated with Packmol-Memgen and subjected to simulations and analyses.**

| System | WT/Mutant | Membrane composition[a] | FFA | Number of FFA molecules in the periplasm | Replicas |
|---|---|---|---|---|---|
| d-PlaF (I) | WT | DOPE:DOPG:C10//DOPE:DOPG 3:1:1//3:1 | C10 | / | 12 |
| d-PlaF (II) | WT | DOPE:DOPG:C10//DOPE:DOPG 3:1:2//3:1 | C10 | / | 12 |
| s-PlaF (I) | WT | DOPE:DOPG:C10//DOPE:DOPG 3:1:1//3:1 | C10 | / | 12 |
| s-PlaF (II) | WT | DOPE:DOPG:C14//DOPE:DOPG 3:1:1//3:1 | C14 | / | 12 |
| t-PlaF (I) | WT | DOPE:DOPG:C10//DOPE:DOPG 3:1:1//3:1 | C10 | / | 12 |
| t-PlaF (II) | WT | DOPE:DOPG:C14//DOPE:DOPG 3:1:1//3:1 | C14 | / | 12 |
| t-PlaF (III) | WT | DOPE:DOPG 3:1 | C10 | 10 | 12 |
| t-PlaF (IV) | F229W | DOPE:DOPG 3:1 | C10 | 9 | 12 |
| t-PlaF (V) | L177W | DOPE:DOPG 3:1 | C10 | 10 | 12 |

[a] Composition of the “upper” // “lower” leaflet; the “upper” leaflet is oriented to the periplasmic space.

**TABLE S4. Thermal stability measured by nanoDSF.**

| Protein | $T_m$/°C |
|---|---|
| PlaF | 55.6 |
| PlaF$_{R187A}$ | 50.3 |
| PlaF$_{E216A}$ | 50.2 |
| PlaF$_{R187A\text{-}A220G}$ | 51.8 |
| PlaF$_{E216A\text{-}A220G}$ | 52.2 |
| PlaF$_{R217A}$ | 52.8 |
| PlaF$_{R187A\text{-}E216A}$ | 48.0 |

**TABLE S5. Pulling points across the T3 for sMD simulations.**

| **T3 pulling points [a]** | **Amino acid residues** |
|---|---|
| 0[b] | S137 |
| I | S137, M138, L184 |
| II | F174, F229, L232 |
| III | K170, Q234, Y236 |
| IV | K170, Q234, Y236 |

[a] Pulling points are COM of backbone atoms of corresponding amino acid residues.

[b] The OH group of S137 was considered the pulling point.

**TABLE S6. Structural stability of proposed T3 tunnel variants of PlaF determined using FoldX and corresponding influence on tunnel characteristics calculated with CAVER.**

| PlaF variant | $\Delta\Delta G$[a,b] | Average bottleneck radius[c,d] | Average length[d] |
|---|---|---|---|
| L177W | 0.53 | 1.70 | 18.51 |
| F229W | 0.81 | 1.67 | 17.77 |

[a] $\Delta\Delta G = \Delta G_{\text{variant}} - \Delta G_{\text{wild type}}$.

[b] In kcal mol$^{-1}$.

[c] Data calculated with a probe radius of 1.2 Å.

[d] In Å.

### Supplementary figures

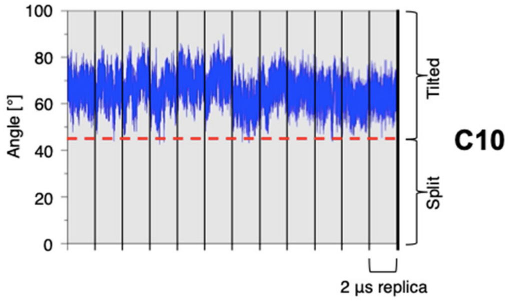

**Figure S1. Unbiased MD simulations of monomeric PlaF in the presence of FFA in the upper leaflet (DOPE:DOPG:C10 of 3:1:1).** When starting from t-PlaF, in all replicas the structure remains tilted after 2 μs.

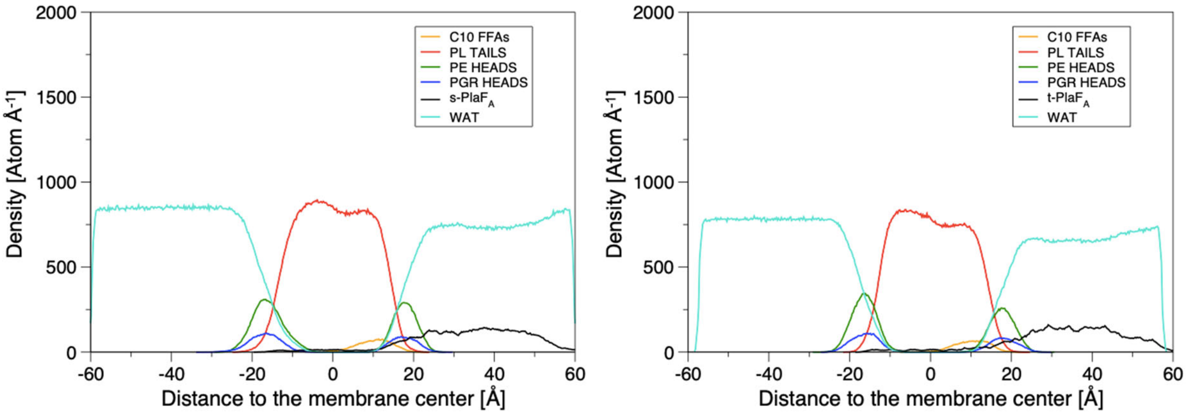

**Figure S2. Atom density profiles of membrane components averaged over 12 independent, unbiased MD simulations of s-PlaF and t-PlaF configurations** for the phospholipid (PL) tails and head groups (PE and PGR) and decanoic acid (C10) molecules. The obtained shapes correspond with those generally found by experiments and MD simulations for biomembranes and our previous work (1, 6-8). Both plots present the composition of the upper leaflet of DOPE:DOPG:C10 3:1:1.

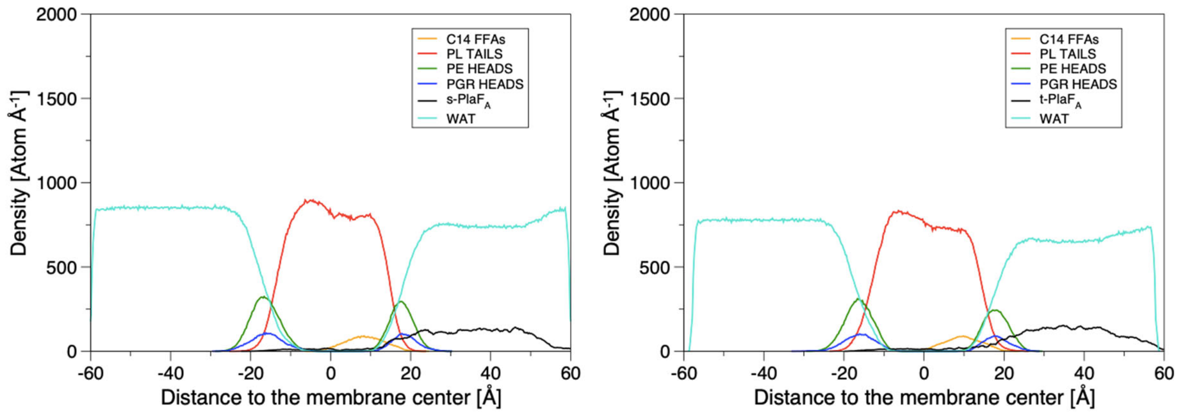

**Figure S3. Atom density profiles of membrane components averaged over 12 independent, unbiased MD simulations of s-PlaF (left) and t-PlaF (right) configurations** for the phospholipid (PL) tails and head groups (PE and PGR) and myristic acid (C14) molecules. The obtained shapes correspond with those generally found by experiments and MD simulations for biomembranes and our previous work (1, 6-8). Both plots present the composition of the upper leaflet of DOPE:DOPG:C14 3:1:1.

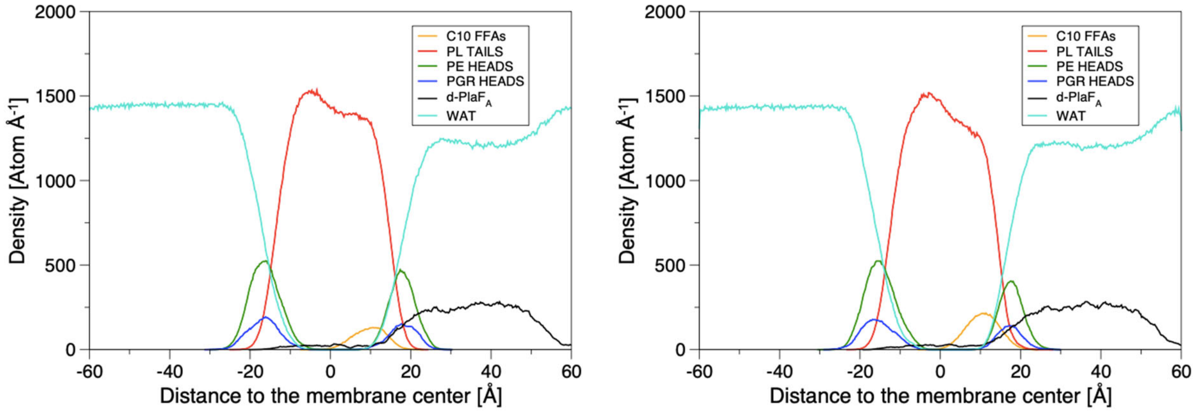

**Figure S4. Atom density profiles of membrane components averaged over 12 independent, unbiased MD simulations of d-PlaF configurations** for the phospholipid (PL) tails and head groups (PE and PGR) and decanoic acid (C10) molecules. The obtained shapes correspond with those generally found by experiments and MD simulations for biomembranes and our previous work (1, 6-8). The left plot corresponds to the composition of the upper leaflet of DOPE:DOPG:C10 3:1:1. The right plot corresponds to the composition of the lower leaflet of DOPE:DOPG:C10 3:1:2.

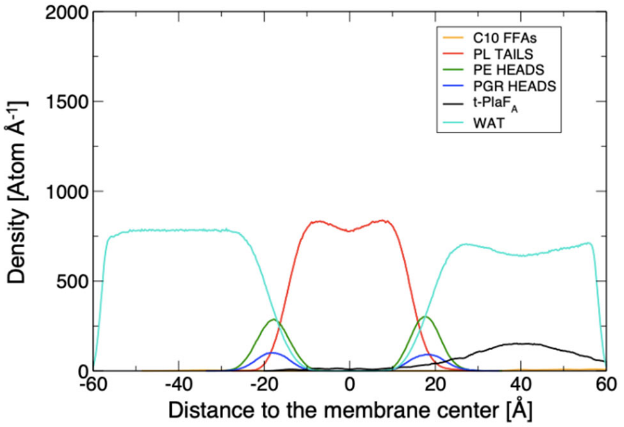

**Figure S5. Atom density profiles of membrane components averaged over 12 independent, unbiased MD simulations of t-PlaF configurations** for the phospholipid (PL) tails and head groups (PE and PGR) and decanoic acid (C10) molecules. The obtained shapes correspond with those generally found by experiments and MD simulations for biomembranes and our previous work (1, 6-8). The plot corresponds to a membrane composition of DOPE:DOPG 3:1. 30 mM decanoic acid is added to the periplasmic space soluble fraction (10 molecules).

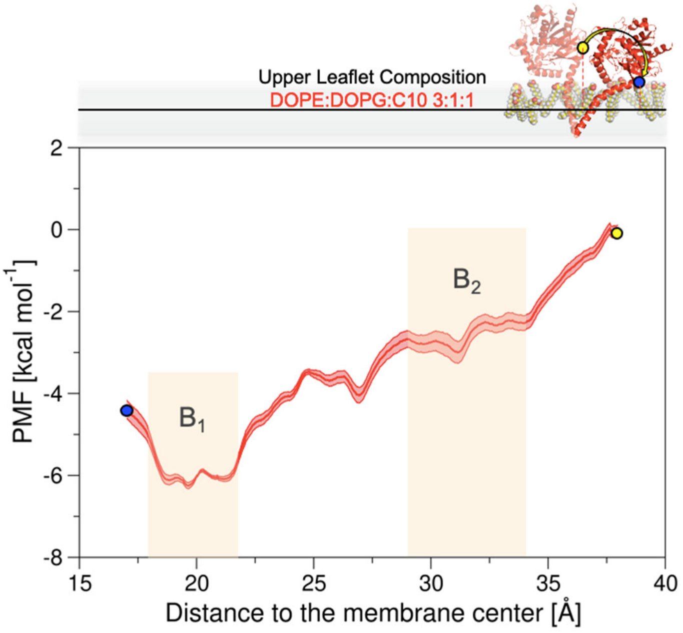

**Figure S6. PMF of monomer tilting in the presence of fatty acids in the membrane (system s-PlaF(I)).** The distance between the COM of $C_\alpha$ atoms of residues 33-37 (blue and yellow spheres) and the COM of the $C_{18}$ of the oleic acid moieties of all lipids in the membrane (continuous horizontal line in the membrane slab) was used as a reaction coordinate. The shaded area shows the standard error of the mean across the entire profile. The orange-shaded regions are the integration limits used to calculate $K_{tilting}$ (eq. S5). The spheres in the PMF relate to the monomer configurations shown in the inset.

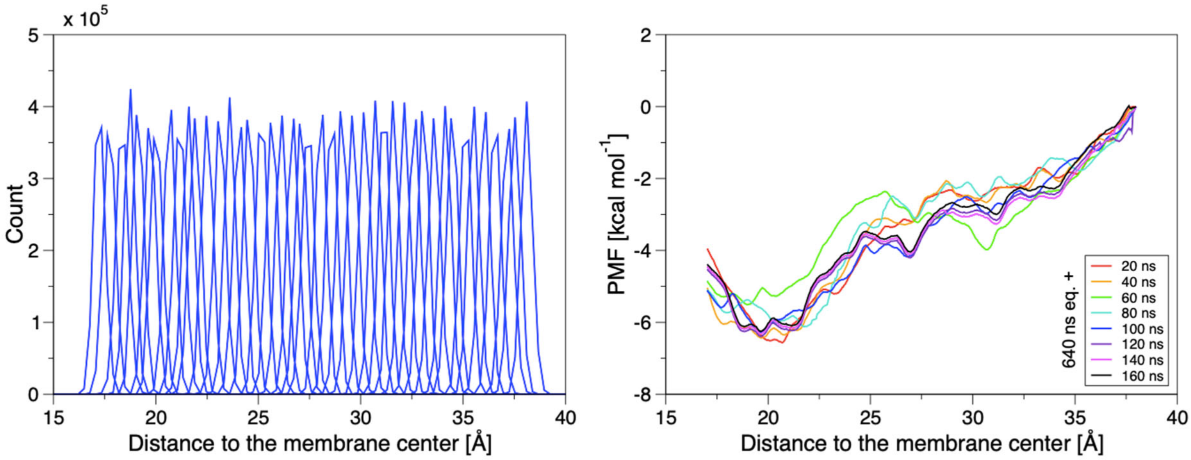

**Figure S7. Evaluation of the PMF of monomer tilting in the presence of fatty acids in the membrane (system s-PlaF(I)). A)** Histograms indicate sufficient overlap among the umbrella windows using a force constant of 4 kcal $mol^{-1}$ $Å^{-2}$: the minimum window overlap is 23.5% (mean ± SEM: 34.9 ± 1.4%) between contiguous windows. **B)** Succession of PMFs with increased sampling times per window after 640 ns of equilibration time. The plot indicates a converged PMF after 800 ns of sampling.

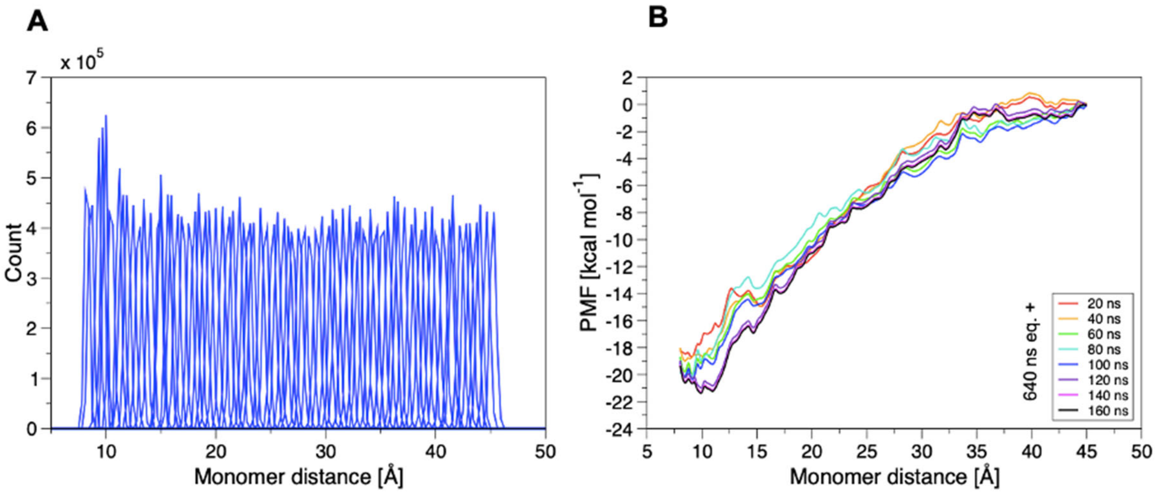

**Figure S8. Evaluation of the PMF of dimer separation in the presence of fatty acids into the membrane (system d-PlaF (I)). A)** Histograms indicate sufficient overlap among the umbrella windows using a force constant of 4 kcal $mol^{-1}$ $Å^{-2}$, the minimum window overlap is 7.9% (mean ± SEM 32.2 ± 1.5%) between contiguous windows. **B)** Succession of PMFs with increased sampling times per window after 640 ns of equilibration time. The plot indicates a converged PMF after 800 ns of sampling.

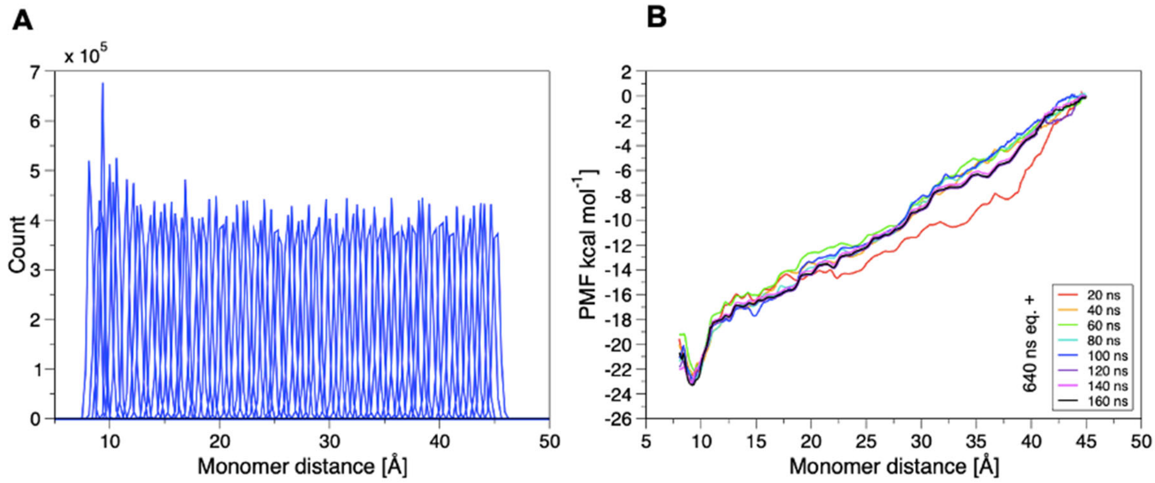

**Figure S9. Evaluation of the PMF of dimer separation in the presence of fatty acids into the membrane (system d-PlaF (II)). A)** Histograms indicate sufficient overlap among the umbrella windows using a force constant of 4 kcal mol$^{-1}$ Å$^{-2}$, the minimum window overlap is 14.2% (mean ± SEM 33.9 ± 1.4%) between contiguous windows. On the right, the convergence plot indicates sufficient sampling time for the selected system. **B)** Succession of PMFs with increased sampling times per window after 640 ns of equilibration time. The plot indicates a converged PMF after 800 ns of sampling.

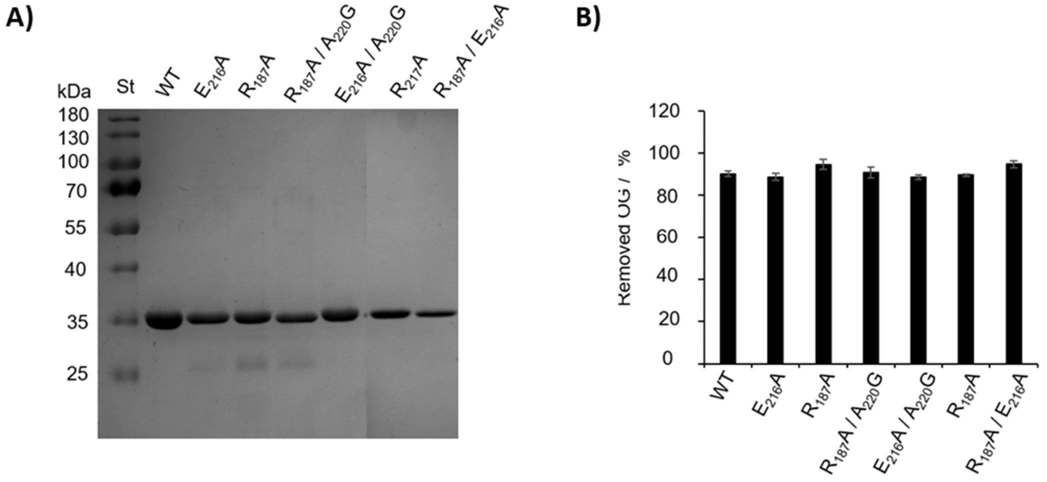

**Figure S10: Purification and removal of OG from reconstituted $PlaF_{WT}$ and LL domain variants.** **A)** PlaF and LL domain variants were purified in the presence of 20 mM OG and analyzed by SDS-PAGE (12%). Molecular weights of protein standard in kDa are indicated. **B)** OG is removed from the reconstituted PlaF samples after 1 h incubation with BioBeads. The results are mean ± SD of two experiments.

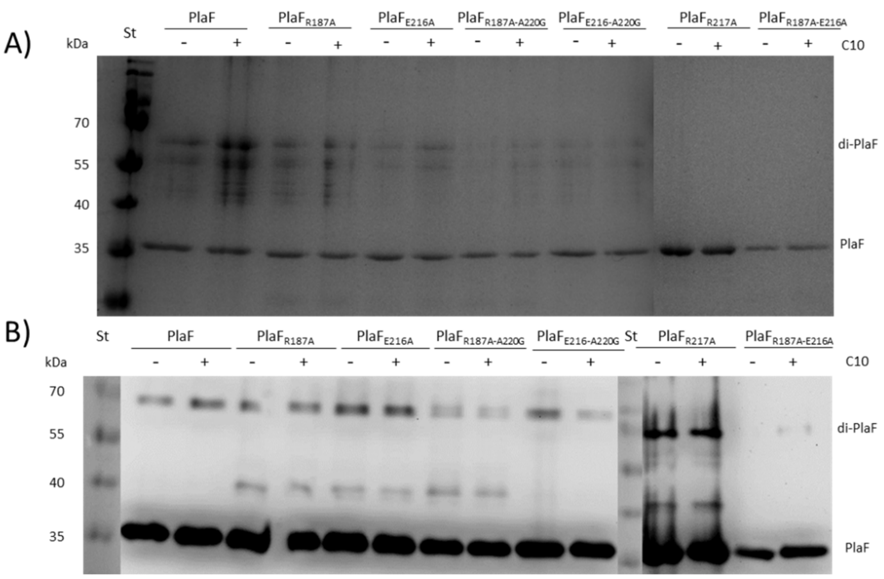

**Figure S11: The effect of C10 on PlaF dimerization. A)** SDS-PAGE and **B)** western blot analyses of PlaF and lid-like domain protein variants reconstituted into DOPE:DOPG SUV and incubated with 150 mM DMP crosslinker in the presence or absence of 12 mM FA C10 for 2 h at room temperature. The same amounts of each protein were loaded onto SDS-PAGE.

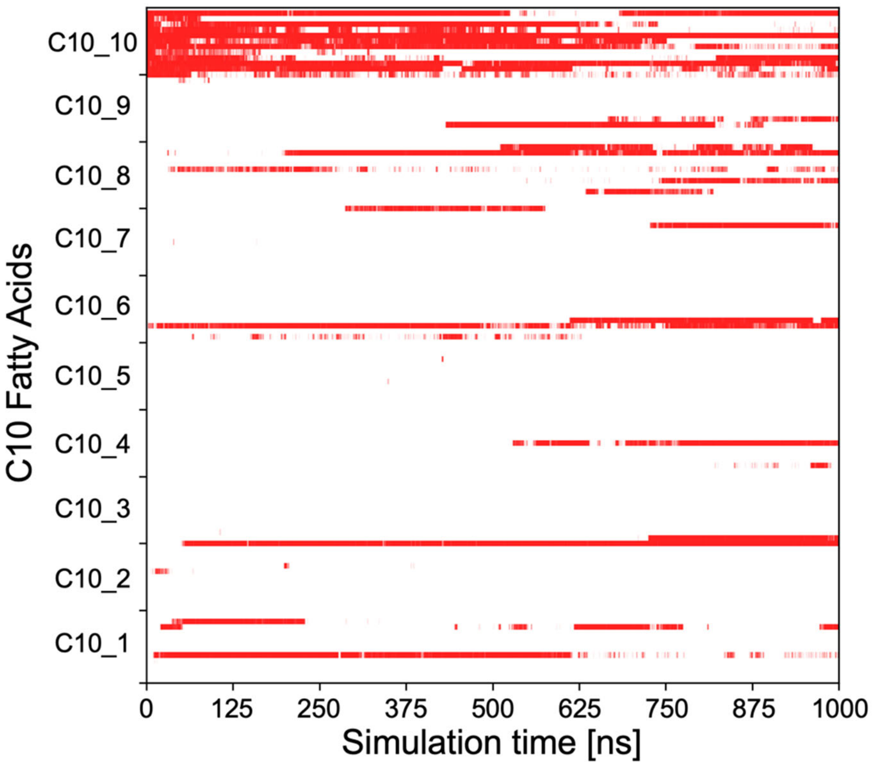

**Figure S12. Free ligand diffusion simulations of 10 C10 around PlaF (System t-PlaF (III)).** Horizontal consecutive points indicate the long residence of fatty acid molecules with a distance $d < 5$ Å with respect to the T3 entrance. $d$ is calculated between the COM of fatty acid atoms and the COM of residues K170, Q234, and Y236 considered the T3 entrance.

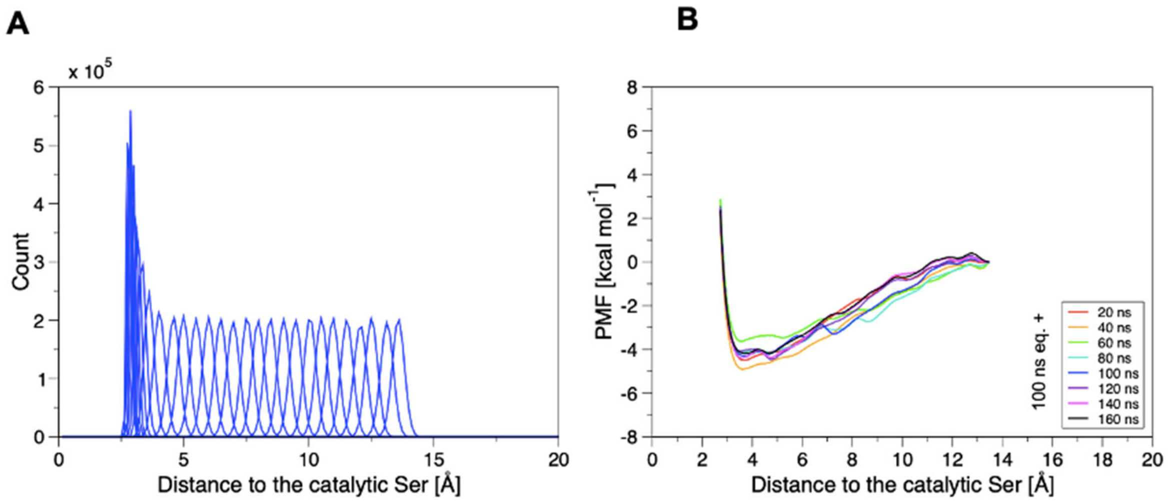

**Figure S13. Evaluation of the PMF of fatty acid-T3 extraction (System t-PlaF (III)). A)** Histograms indicate sufficient overlap among the umbrella windows using a force constant of 4 kcal $mol^{-1}$ $Å^{-2}$; the minimum window overlap is 21.3% (mean ± SEM 33.6 ± 3.1%) between contiguous windows. **B)** Succession of PMFs with increased sampling times per window after 100 ns of equilibration time. The plot indicates a converged PMF after 260 ns of sampling

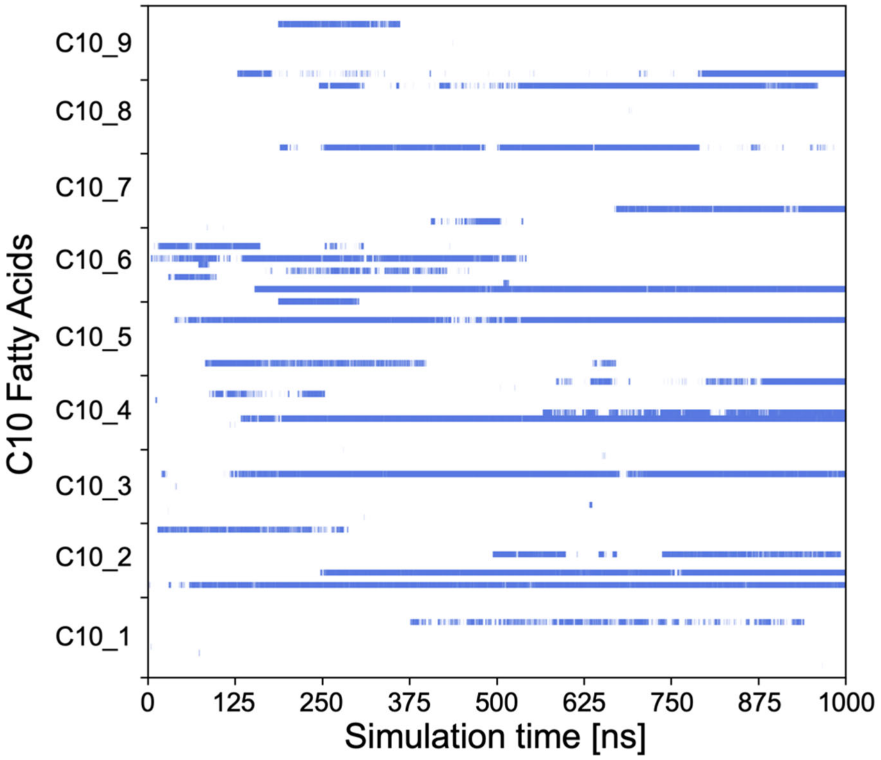

**Figure S14. Free ligand diffusion simulations of 10 C10 around PlaF (System t-PlaF (IV)).** Horizontal consecutive points indicate the long residence of fatty acid molecules with a distance $d < 5$ Å with respect to the T3 entrance. $d$ is calculated between the COM of fatty acid atoms and the COM of residues K170, Q234, and Y236 considered the T3 entrance.

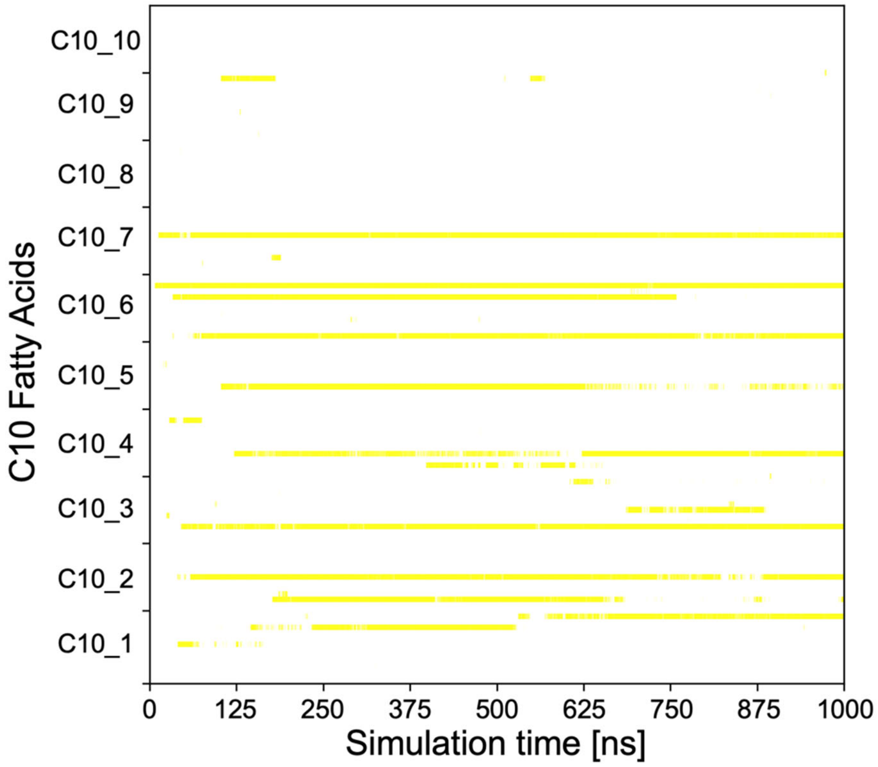

**Figure S15. Free ligand diffusion simulations of 10 C10 around PlaF (System t-PlaF (V)).** Horizontal consecutive points indicate the long residence of fatty acid molecules with a distance $d < 5$ Å with respect to the T3 entrance. *d* is calculated between the COM of fatty acid atoms and the COM of residues K170, Q234, and Y236 considered the T3 entrance.

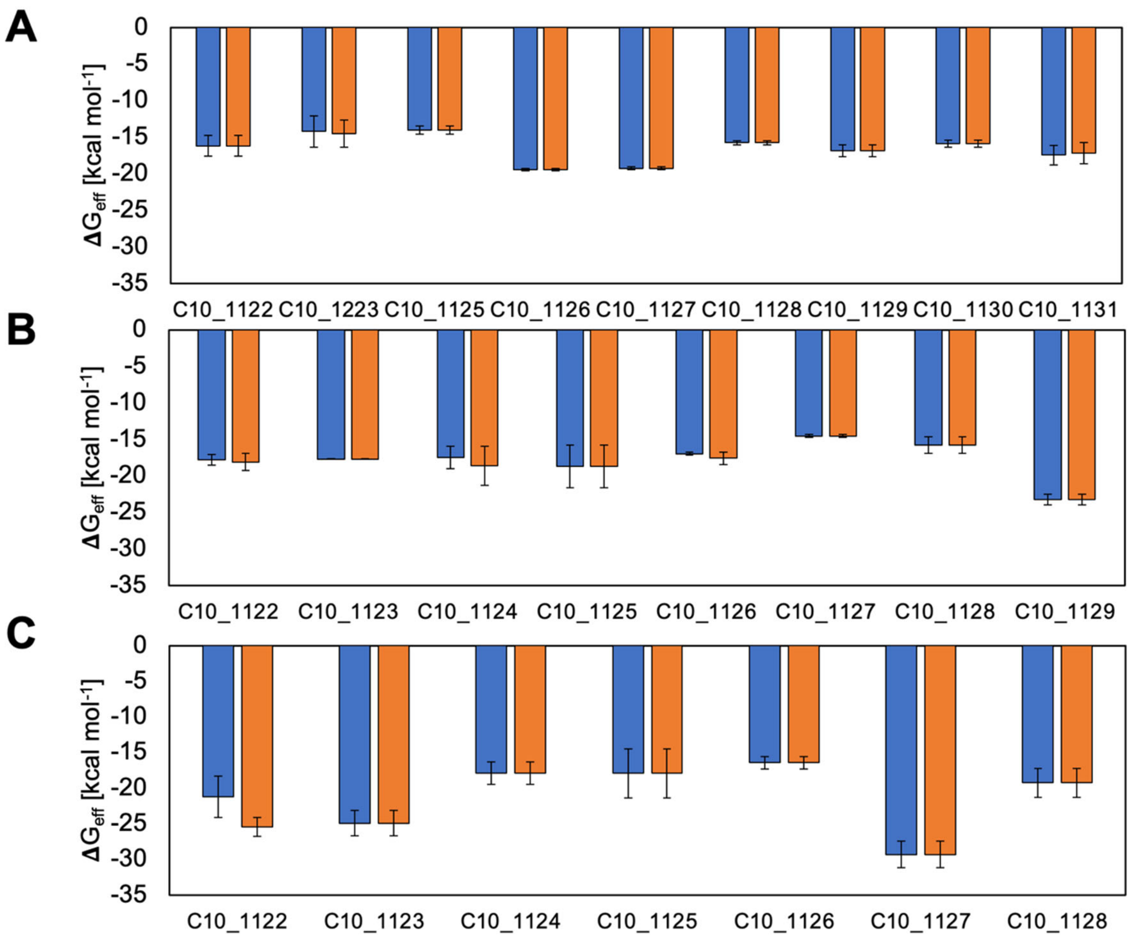

**Figure S16. MM-PBSA binding effective energy computations of C10 to WT and single mutants in T3 are converged. A) WT.** Blue bars describe the binding effective energy from the first half of the replica averaged over the five top-ranked clusters for each of the nine fatty acids. Orange histograms describe the results for the second half of the replica. The error bars represent the SEM. **B) F229W.** Blue bars describe the binding effective energy from the first half of the replica averaged over the five top-ranked clusters for each of the eight fatty acids. Orange histograms describe the results for the second half of the replica. The error bars represent the SEM. **C) L177W**. Blue bars describe the binding effective energy from the first half of the replica averaged over the five top-ranked clusters for each of the seven fatty acids. Orange histograms describe the results for the second half of the replica. The error bars represent the SEM.

**Supplementary movies**

**Movie S1: Extraction of a C10 fatty acid molecule via the tail from the bound position inside T3 of t-PlaF using sMD simulations.** The grey cartoon represents t-PlaF, which is embedded in a membrane composed of DOPE:DOPG 3:1. At the start of the simulation, the fatty acid tail reaches close to the catalytic S137 (purple sticks).
